# Ecological causes of uneven mammal diversity

**DOI:** 10.1101/504803

**Authors:** Nathan S. Upham, Jacob A. Esselstyn, Walter Jetz

## Abstract

The uneven distributions of species over geography (e.g., tropical versus temperate regions) and phylogeny (e.g., rodents and bats versus the aardvark) are prominent biological patterns for which causal interconnections remain enigmatic. Here we investigate this central issue for living mammals using time-sliced clades sampled from a comprehensive recent phylogeny (*N*=5,911 species, ∼70% with DNA) to assess how different levels of unsampled extinction impact the inferred causes of species richness variation. Speciation rates are found to strongly exceed crown age as a predictor of clade species richness at every time slice, rejecting a clock-like model in which the oldest clades are the most speciose. Instead, mammals that are low-vagility or daytime-active show the fastest recent speciation and greatest extant richness. This suggests primary roles for dispersal limitation leading to geographic speciation (peripatric isolation) and diurnal adaptations leading to ecological speciation (time partitioning). Rates of speciation are also faster in temperate than tropical lineages, but only among older clades, consistent with the idea that many temperate lineages are ephemeral. These insights, enabled by our analytical framework, offer straightforward support for ecological effects on speciation-rate variation among clades as the primary cause of uneven phylogenetic richness patterns.

## INTRODUCTION

Biological diversity is concentrated more at the equator than the poles, and more in some clades than others. Yet whether the latitudinal pattern of variation causes the phylogenetic one is an open question. The latitudinal diversity gradient is generally attributed to tropical biomes being stable, productive, and old (Fine & Ree 2006; Jablonski *et al*. 2006; Mittelbach *et al*. 2007; Jetz & Fine 2012; Jansson *et al*. 2013; Pontarp *et al*. 2019), but there is less consensus regarding why species richness is distributed unevenly across the tree of life. Phylogenetic tree shape was first characterized taxonomically (Willis 1922) and later formalized under the concept of tree imbalance or unevenness (Mooers & Heard 1997). To arise, more speciose clades must derive from faster net diversification (speciation – extinction), older ages (earlier divergences), or both. However, the relative contribution of clade rates and ages to species richness is widely disputed (e.g., (McPeek & Brown 2007; Wiens 2011; Rabosky *et al*. 2012; Hedges *et al*. 2015)). Empirical phylogenies might record diversification-rate variation due to stochastic factors, determinism (e.g., via ecological factors), or artifacts of how we reconstruct evolutionary history (Ricklefs 2003; Blum & François 2006; Phillimore & Price 2008; Rabosky 2009; Venditti *et al*. 2010; Davies *et al*. 2011; Purvis *et al*. 2011; Price *et al*. 2012; Moen & Morlon 2014; Castro-Insua *et al*. 2018; Machac *et al*. 2018; Diaz *et al*. 2019; Louca & Pennell 2020). Latitude might alter the rates at which new species originate, persist, or go extinct (Jablonski *et al*. 2006; Weir & Schluter 2007; Cutter & Gray 2016; Machac & Graham 2017; Silvestro *et al*. 2020), but so too might species’ intrinsic traits (Jablonski 2008), some of which are correlated with latitude (e.g., (Alroy 2019)). Thus, understanding the processes underpinning uneven species richness requires connecting direct (e.g., rates, ages) and indirect (e.g., ecological) causes to tease apart their joint influences upon the phylogenetic distribution of species richness.

The challenge of disentangling the relative importance of clade ages (time) versus rates of speciation and extinction (whether stochastic or ecologically deterministic) suggests the need to establish a hierarchical framework uniting these direct and indirect potential causes of uneven species richness (Fig. 1). To do so, we propose the following set of hypotheses: H_0_, speciation and extinction rates among clades do not vary substantially, so clade ages best explain uneven species richness; or, H_A_, differences in among-clade diversification rates explain species richness better than clade age. If the alternative hypothesis is supported, then certain ecological attributes (e.g., patterns of space use, activity period, or environment niche) may explain changes in diversification rates, and thereby indirectly cause patterns of uneven species richness (Fig. 1b).

**Fig. 1.**
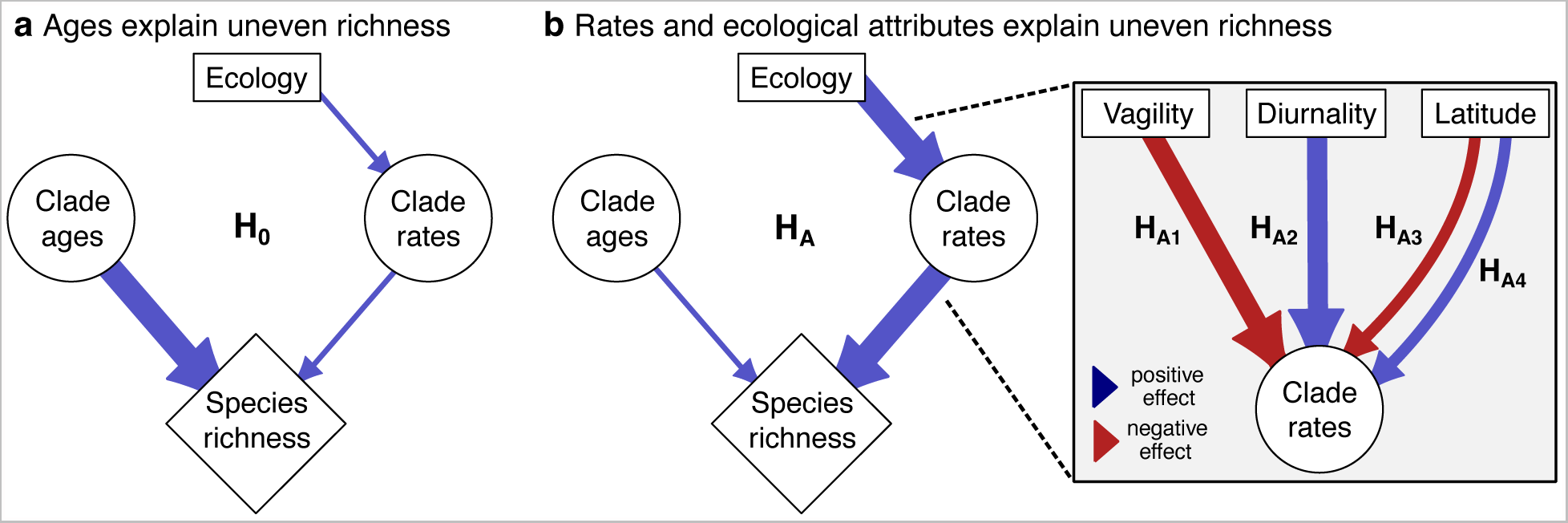
Hypotheses for uneven species richness among branches in the tree of life. The observation that different groups of species (clades) have differing numbers of extant species is generally explained in one of two ways: (a) by uneven clade ages, in which younger clades have few species while older clades are the most speciose (null hypothesis, H_0_); or (b) by uneven macroevolutionary rates (speciation – extinction = net diversification), in which the most speciose clades have the fastest rates (alternative, H_A_). H_A_ implies that some varying ecological attributes of species are, in turn, responsible for the observed variation in clade rates. We specifically assess three aspects of ecology (inset): vagility, a measure of individual dispersal ability in a species (H_A1_); diurnality, or the propensity for individuals in a species to be active during daylight hours (H_A2_); and centroid latitude of a species’ geographic range, a surrogate metric capturing abiotic conditions such as environmental stability (H_A3_ and H_A4_). See the main text for explanations of each attribute’s expected directions of influence upon clade rates.

This framework can be tested using phylogenetic path analysis (von Hardenberg & Gonzalez-Voyer 2013), which unifies macro-scale approaches from ecology and evolution. In macroecology, environmental variables or species traits are often related directly to species richness (e.g., associations between clade richness and mean body size; (Gittleman & Purvis 1998; Isaac *et al*. 2005)). In macroevolution, measurements of clade rates or ages are either compared to species richness (e.g., (Scholl & Wiens 2016; Sánchez-Reyes *et al*. 2017)) or ecological traits (e.g., (Beaulieu & O’Meara 2016; Harvey & Rabosky 2018)), but usually not both. Recently, Harvey et al. (2020) used a path analysis of geographic regions to establish causality between environmental variables, species richness, and speciation rates in suboscine birds, which revealed the spatial context of uneven species diversification. However, the phylogenetic context of why species richness varies so dramatically among clades remains enigmatic, particularly regarding the relative causality of temporal and ecological factors.

A major limitation to studying among-clade diversity has been the reliance, as units of analysis, on higher taxa (e.g., (Rabosky *et al*. 2012; Castro-Insua *et al*. 2018)), which often have vast differences in age. For example, crown ages of mammal families range from 3.8 to 59.0 million years (Ma; (Upham *et al*. 2019)). To avoid comparing heterogeneously defined clades, we propose an objective strategy for delimiting analytically equivalent clades, i.e., ‘time-sliced clades.’ By slicing a phylogeny at a given time, we can then take the tipward groups as objective units of analysis. As shown for an example family of mammals (Fig. 2), slicing the tree at 5 Ma results in the delimitation of 17 clades containing two or more species. Those clades are united by having (i) a stem age >5 Ma and crown age < 5 Ma; and (ii) a rootward backbone of shared evolutionary history. However, because each clade varies in estimated crown age, species richness, clade summaries of species attributes, and species-specific ‘tip’ speciation rates, there is a key opportunity for identifying partial associations (Fig. 2c). Critically, defining time-sliced clades at successively older time-points within a phylogeny of extant species yields the expectation for progressively greater bias from unsampled extinction events — i.e., Marshall’s (2017) ‘fifth law of palaeobiology’ (Fig. 2a). As a result, speciation rates will be increasingly underestimated or ‘pulled’ from the actual rates of the birth-death process in older clades (Kubo & Iwasa 1995; Louca & Pennell 2020). This general expectation has been confirmed in crown Mammalia, for which fossil- and molecular-based speciation rates overlap only between 0 and ∼10 Ma (Upham *et al*. 2021). Thus, analyzing clades from shallow to deep time slices of an extant phylogeny presents a further opportunity for assessing the impact of unsampled extinctions on among-clade correlations in species attributes, rates, ages, and diversities.

**Fig. 2.**
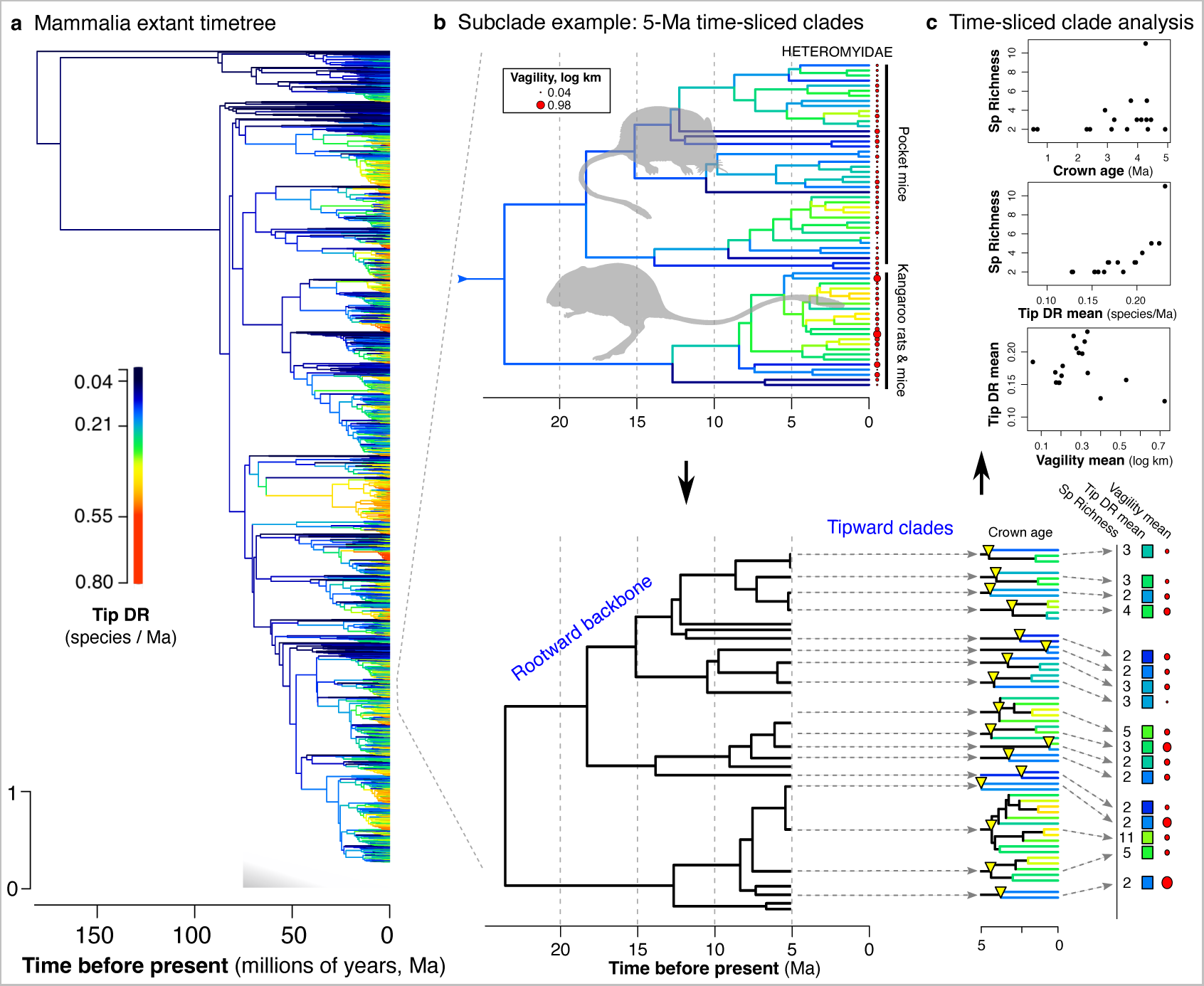
Approach of using time-sliced clades to test eco-evolutionary hypotheses. (a) The mammal timetree is painted with species-specific (tip) speciation rates calculated using the tip DR metric across the full tree. The fraction of unsampled extinction events is expected to increase at deeper levels of the extant timetree, with only two surviving lineages (leading to extant therians and monotremes) sampled at the root of crown Mammalia. (b) Example of how a subclade of mammals can be divided into time-sliced clades, here the rodent family Heteromyidae (64 species) with clades delimited tipward of an arbitrary line drawn at 5-million years (Ma; branch colors correspond to tip DR). (c) The crown age of those clades with two or more species are by definition < 5 Ma (yellow triangles), and the species richness values range from 1-11 species in this example. Also summarized are the clade harmonic mean of tip DR (tip DR mean), and the clade geometric mean of vagility values (or other ecological traits). The rootward backbone of those time-sliced clades represents their expected covariance structure for use in comparative analyses (e.g., phylogenetic generalized least squares). Analyses conducted in Fig. 4 and 5 are based upon summary values for time-sliced clades delimited in this manner.

Here, we apply this novel clade-level framework to the investigation of temporal and ecological causes of uneven diversification in Mammalia. Considering the ∼6,500 recognized living species of mammals (Burgin *et al*. 2018; MDD 2023), we see that similarly aged clades range from mega-diverse rodents (∼2,600 living species) and bats (∼1,400 species) to species-poor groups like treeshrews (23 species) and pangolins (8 species, all four clades share stem ages of ∼60-70 Ma; (Meredith *et al*. 2011; Upham *et al*. 2019; Álvarez-Carretero *et al*. 2022; Foley *et al*. 2023)). We here focus on three ecological factors hypothesized to influence rates of mammal speciation — vagility, diurnality, and latitude (Fig. 1b) — as measured on extant time-calibrated phylogenies (i.e., timetree). First, we tested whether low-vagility species have faster speciation than more dispersive species given their greater likelihood of forming peripheral isolates (H_A1_) (Mayr 1963; Kisel & Barraclough 2010). For this test, we developed an allopatric index of organismal vagility for all mammals (i.e., maximum natal dispersal distance; (Whitmee & Orme 2013)). Vagility effects have never been assessed across all mammals, although evidence in birds (e.g., (Belliure *et al*. 2000; Claramunt *et al*. 2012)) and reef fishes (Donati *et al*. 2019) supports an inverse vagility-to-speciation rate relationship. Second, we tested whether clades with greater diurnality have increased speciation rates relative to nocturnal clades, following evidence that mammalian ancestors were likely nocturnal until daytime niches evolved ∼35 Ma (H_A2_) (Gerkema *et al*. 2013; Maor *et al*. 2017). A positive influence of diurnality on speciation rates has been found across major tetrapod lineages (Anderson & Wiens 2017), and in primates specifically (Magnuson-Ford & Otto 2012; Santini *et al*. 2015), but has yet to be investigated at the species-level in all mammals (only ancestral diel states have been examined (Maor *et al*. 2017)). Lastly, we examine the effects of latitude on speciation rates, which could either have a negative or positive association (fastest rates at low latitudes, H_A3_, or high latitudes, H_A4_, respectively). Previous work in mammals has supported faster tropical than temperate speciation among orders (Rolland *et al*. 2014), but has been inconclusive among genera (Soria-Carrasco & Castresana 2012) and found the opposite pattern among sister species (Weir & Schluter 2007). Faster temperate than tropical speciation contrasts with the observed pattern of peak tropical mammal richness, meaning that H_A4_ additionally implies higher temperate rates of extinction and thus species turnover (extinction / speciation). This type of ‘ephemeral speciation’ (Rosenblum *et al*. 2012) is supported by observations of faster tip speciation in high-latitude marine fishes (Rabosky *et al*. 2018) and rosid angiosperms (Sun *et al*. 2020), as well as high-elevation birds (Quintero & Jetz 2018), but this hypothesis is so far untested in mammals (reviewed in (Cutter & Gray 2016; Schluter & Pennell 2017)). Drawing upon a comprehensive time-calibrated phylogeny of mammals and tip rates of speciation calculated across a credible set of 10,000 trees (Upham *et al*. 2019), we assembled a corresponding set of species-level ecological traits to query whether factors predicted to cause newly formed species to persist or go extinct are, in turn, causing the observed patterns of uneven species richness among clades.

## METHODS

### Mammalian phylogeny and species trait data

We used the species-level mammal trees of Upham et al. (2019) to conduct all analyses. Briefly, these phylogenies include 5,804 extant and 107 recently extinct species in credible sets of 10,000 trees. They were built using a ‘backbone-and-patch’ framework that applies two stages of Bayesian inference to integrate age and topological uncertainty, and incorporates 1,813 DNA-lacking species using probabilistic constraints (available at <u>vertlife.org/phylosubsets</u>). We compared credible sets of trees built using node-dated backbones (17 fossil calibrations) and tip-dated backbones (matrix of modern and Mesozoic mammals), as well as taxonomically completed trees (5,911 species) versus DNA-only trees (4,098 species) without topology constraints. We calculated phylogenetic signal and tree imbalance statistics using maximum clade credibility (MCC) consensus trees and the R packages “phytools” (Revell 2012) and “apTreeshape” (Bortolussi *et al*. 2006), respectively.

Our workflow for gathering trait data involved (i) unifying multiple trait taxonomies (e.g., EltonTraits v1.0 (Wilman *et al*. 2014), PanTHERIA (Jones *et al*. 2009)) to our phylogeny’s master taxonomy; and (ii) interpolating home range area and vagility to the species level using known allometric relationships in mammals (Fig. S1). Vagility was interpolated as an index value for each species following our updated version of Whitmee and Orme’s (2013) best-fit equation, which applies species means of body mass, home range, and geographic range to calculate the maximum natal dispersal distance per individual (km; Fig. S2). Note that our vagility index does not account for locomotor abilities (e.g., flying or arboreality), but rather captures aspects of space use that scale allometrically across mammals. Collinearity among trait variables was examined using the “corrplot” package in R (Wei 2017).

We identified three species-level traits that are directly related to core hypotheses of ecological diversification while also having minimal collinearity (Fig. S3). These traits are: (i) an allometric index of vagility based on a best-fitting equation of log(maximum natal dispersal distance, km) = -2.496 + 0.206 log(body mass, g) + 0.323 log(home range size, km) + 0.216 log(geographic range size, km); (ii) diurnality as a binary trait of 0 = nocturnal / cathemeral / crepuscular, and 1 = diurnal; and (iii) latitude calculated as the absolute value of the centroid of a species’ expert geographic range map. As expected (Jetz *et al*. 2004; Whitmee & Orme 2013), the vagility index captures a multivariate signal of average individual space use per species (correlation of r=0.5–0.8 with its component variables; Fig. S3). Diurnality and latitude inform two additional ecological axes, each having low collinearity with vagility or other variables (maximum r=0.34 and 0.13, respectively).

### Tip-level speciation rates

We calculated per-species estimates of expected pure-birth diversification rates for the instantaneous present moment (tips of the tree) using the inverse of the equal splits measure (Steel & Mooers 2010; Jetz *et al*. 2012). This metric has been called ‘tip-level diversification rate’ (tip DR) because it measures recent diversification processes among extant species (Quintero & Jetz 2018). However, to avoid confusion with ‘net diversification’, for which tip DR is not suited when extinction is very high (relative extinction >0.8 (Title & Rabosky 2019)), we here use tip DR as a tip-level speciation rate metric. At the tip level, we confirm that tip DR is tightly associated with model-based estimators of tip speciation and tip net diversification rates in the mammal trees (Fig. S4). At the clade level, Upham *et al*. (2021) showed that (i) the clade-level harmonic mean of tip speciation rates, as measured by tip DR (here called ‘tip DR mean’) approximates the pulled speciation rate for that clade at the instantaneous present (λ_0_), which is an identifiable value (Louca & Pennell 2020); and (ii) the skewness of tip DR in a clade (here called ‘tip DR skew’) approximates that clade’s extent of past diversification-rate shifts, as measured using BAMM rate-shift factors (Rabosky 2014). Thus, we expect tip DR mean and skew to illuminate the speed and heterogeneity of past speciation in a clade, respectively. A critical caveat here, following the demonstration in fig. 4b of Upham *et al*. (2021), is that we only expect the most recent ∼10 Ma of branching in the extant mammal timetree to carry a reliable signal of past speciation-rate dynamics as compared to the fossil record.

**Fig. 3.**
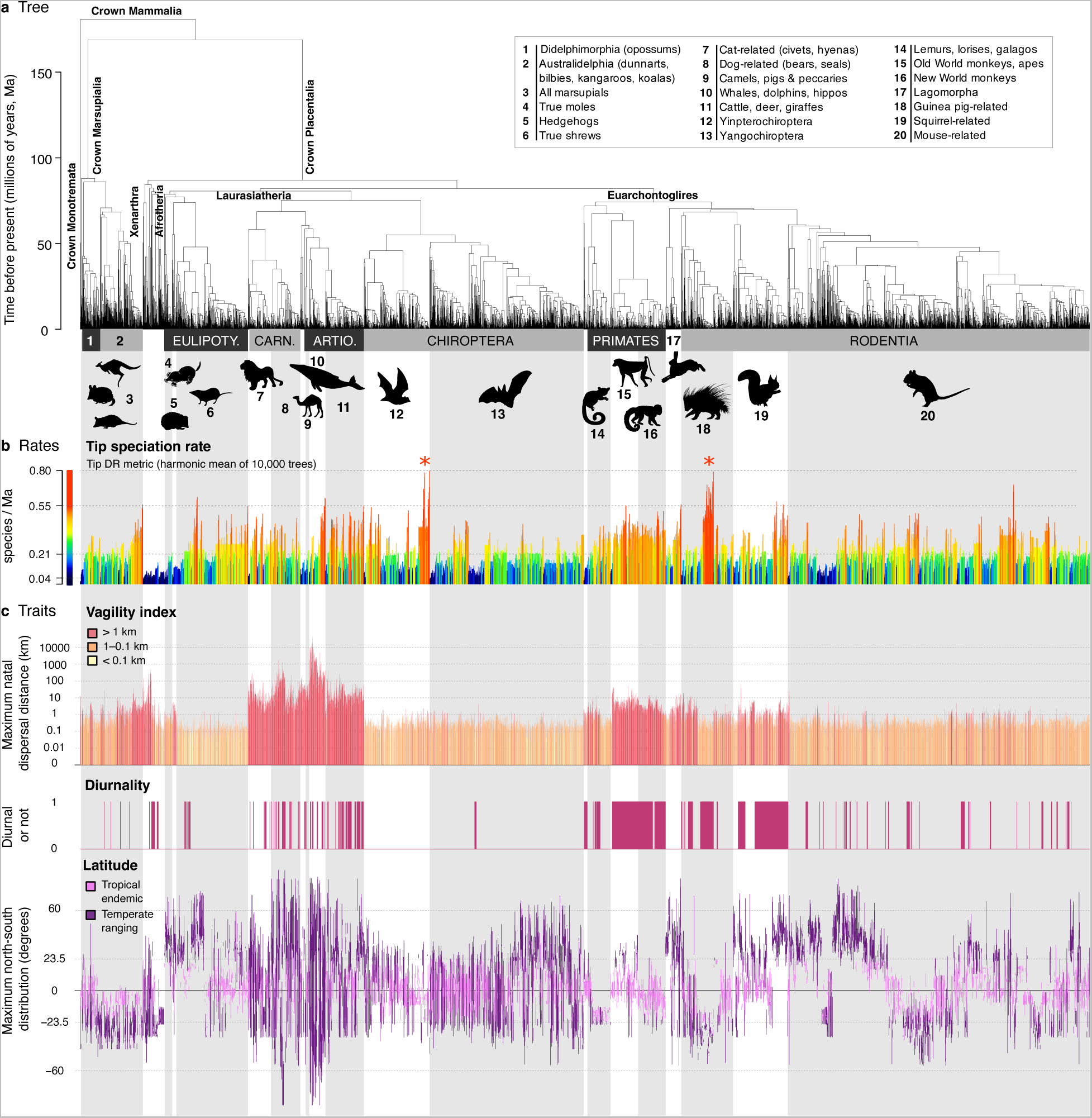
Species-level relationships, rates, and traits for 5,911 living species of mammals globally. (**a**) The maximum clade credibility topology of 10,000 node-dated timetrees, with numbered clade labels corresponding to orders and subclades listed in the plot periphery: Eulipoty., Eulipotyphla; Carn., Carnivora; Artio., Artiodactyla. Scale in millions of years, Ma. (**b**) Tip speciation rates, as measured using tip DR values that correspond to the expected rate of species formation at the instantaneous present for each species (red asterisks refer to the bat clade *Pteropus,* left, and the rodent clade *Ctenomys,* right, which display significantly elevated rates of recent speciation). (**c**) Per-species ecological attributes: allometric index of vagility (dispersal ability), diurnality (predominant daytime activity), and north-to-south latitudinal extent of geographic range map. Silhouettes are from phylopic.org and open-source fonts.

**Fig. 4.**
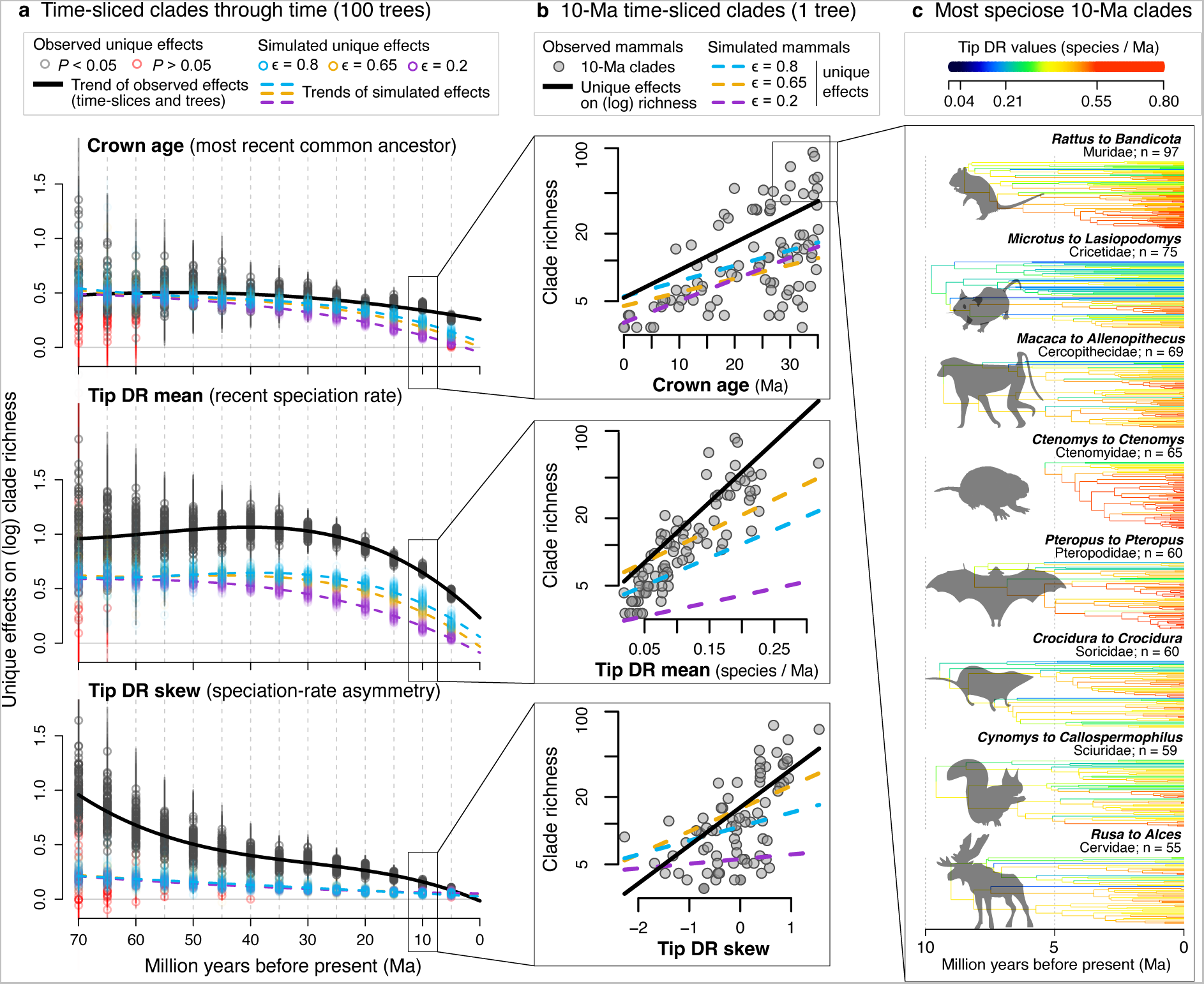
Temporal causes of species richness variation across time-sliced clades. (**a**) Phylogenetic generalized least squares (PGLS) analyses of tipward time-sliced clades (delimited as in Fig. 2) at 5-million-year (Ma) intervals from 5–70 Ma. Analyses are repeated across 100 mammal trees and compared to analogous results from trees simulated under a model of rate-constant birth-death (RCBD). For all clades, partial unique effects are examined on clade richness (PGLS of log clade species richness ∼ crown age + tip DR mean + tip DR skew; predictors are standardized). Tip DR mean has consistently stronger unique effects than does crown age, and stronger than expected from the stochastic rate variation in RCBD simulations (95% confidence intervals do not overlap from 5–30 Ma). At deeper time slices, tip DR skew also explains more variation in clade richness than expected from simulations. (**b**) Each gray point in part **a** represents the unique-effect PGLS slope (solid black line) for a set of mammal time-sliced clades, here shown for an example of 10-Ma clades in 1 mammal tree compared to analogous clades from RCBD simulations (dotted colored lines under different extinction fractions, ε). Note the clade-level predictors are shown with original (non-standardized) values for illustration purposes only. (**c**) Illustration of the most speciose 10-Ma clades delimited from 1 mammal tree (n = species richness of clade). Branches are colored relative to species’ tip DR values (interior nodes painted for visualization purposes using a Brownian motion reconstruction). Silhouettes are from phylopic.org and open-source fonts.

### Time-sliced clades and clade-level tests of species richness variation

To objectively define clades, we arbitrarily drew lines (referred to as “time slices”) at 5-Ma intervals and took the resulting *tipward* (all the way to the extant tip) clades as non-nested units of analysis. The *rootward* relationships of those clades (the “rootward backbone”) was retained for each interval, giving the expected covariance structure among clades when performing phylogenetic generalized least squares (PGLS) analyses (see Fig. 2 for illustration). We used the “treeSlice” function in phytools to construct clade sets across mammal timetrees and three sets of rate-constant birth-death (RCBD) simulated trees. These RCBD simulations were run using the “pbtree” function in phytools under scenarios of extinction fraction, ε, matching the empirical median (ε=0.65) versus low (ε=0.2) or high (ε=0.8) values, and with simulations set to 5,911 species and re-scaled to crown age of 188 Ma to approximate the branching history of extant mammals. We also compared the time-sliced clade results to analyses based on traditional named clades (genera, families, and orders). All PGLS analyses were performed excluding extinct species, using Pagel’s “lambda” transformation in phylolm (optimized for large trees (Ho & Ané 2014)), and repeating the analysis on 100 or 1,000 trees. We also performed multivariate analyses including percent of DNA-sampled species per clade (for the completed trees) to test whether any results were affected by the proportion of taxonomically imputed species.

### Tip-level tests of speciation-rate correlates

To examine correlative structures underlying observed tip-rate variation, we performed tip-level PGLS analyses between species’ ecological traits and tip DR values across 1000 trees, focusing on a 5,675-species data set that excluded all extinct (n=107) and marine (n=129) species. We followed Freckleton et al. (2008) in using trait ∼ rate models in our tip-level PGLS analyses to avoid identical residuals in the dependent variable (i.e., sister species have identical tip DR values, which otherwise violates the assumed within-variable data independence in bivariate normal distributions). The trait ∼ rate approach was previously applied using tip DR in univariate contexts (Harvey *et al*. 2017), and performs well compared to QuaSSE (Harvey & Rabosky 2018).

### Clade-level tests of speciation-rate correlates

At the clade level, univariate PGLS was performed typically (rate ∼ trait models), since clade tip DR mean gave independent values to sister clades. These analyses were conducted on 1,000 trees as above, except that per-clade trait summaries were standardized (mean centered, standard deviation scaled) using geometric means for vagility and arithmetic means otherwise. For focal species attributes, we took clade-level average values to be robust ecological summaries of clades in relation to other clade metrics.

### Phylogenetic path analyses

We performed path analysis aiming to fully resolve correlational structures and thereby translate from the language of statistical association to causality. For phylogenetic path analyses, we used PGLS to test statements of conditional independence (von Hardenberg & Gonzalez-Voyer 2013) across 27 pre-selected path models (Fig. S5). For each tree and clade set, we used the “phylopath” R package (van der Bijl 2018) to analyze models and perform conditional model averaging. Time-sliced clades at 10-, 30-, and 50- Ma intervals were analyzed and compared to somewhat analogous taxon-based clades of genera, families, and orders, with the expectation that older clades will contain greater unmeasured extinction and thus be less reliable indicators of historical speciation rates.

## RESULTS

### Unevenness of traits, rates, and species in the mammal timetree

We find dramatic variation in the phylogenetic distribution of mammal biodiversity (Fig. 3). The shape of the extant phylogeny is highly imbalanced, either as measured on the DNA-only or completed MCC tree (Colless’ *I* values of 53591 and 82550, respectively) as compared to Yule expectations (test statistics of 5.872 and 6.397; *P* < 0.001). For speciation-rate estimates, we similarly find the tip DR values are non-uniform with respect to taxonomic orders (Kruskal-Wallis *χ*^2^ = 1085.3, df = 26, *P* < 0.001) and families (*χ*^2^ = 2564.4, df = 161, *P* < 0.001). Variation among species attributes is also non-uniform, as illustrated by significant phylogenetic signal with respect to vagility (*K* = 0.045, *P* = 0.034 [1,000 randomizations]), diurnality (*K* = 0.137, *P* = 0.001), and latitude (*K* = 0.062, *P* = 0.001; using the DNA-only MCC tree, as imputed species can bias studies of trait evolution, see (Rabosky 2015)).

Visually, mapping rates and traits on the timetree reveals a macroscopic view of global mammal diversification history. Two major pulses of recent speciation are apparent in bats and rodents, corresponding to the largest and most reliable diversification-rate shifts in mammals ((Upham *et al*. 2021); Fig. 3b, red asterisks in clades 12 and 18). We find greater vagility and latitudinal extents within Carnivora and Artiodactyla (clades 7–11; Fig. 3c) as compared to other mammal orders, along with a more heterogenous mix of diurnal or non-diurnal activity. In contrast, simian primates and squirrel-related rodents show clade-wide sweeps of diurnality (clades 15, 16, 19). We also find a conspicuous latitudinal pattern of alternating north-south-north-south endemism from Lagomorpha to Rodentia (clades 17–20), which is difficult to dismiss as randomness given the clade-wise sorting of traits, so it may reflect biogeographic incumbency effects. We also note the clear signal of Madagascar visible from 12 to 24 degrees south latitude, representative of endemic radiations of tenrec afrotherians, euplerid carnivorans, strepsirrhine primates, and nesomyid rodents.

### Effects of ages and rates on clade species richness

To separate the putative temporal causes of among-clade richness differences, we performed PGLS analyses on time-sliced clades. Univariate analyses show that crown age, tip DR mean, and tip DR skew are consistently the best predictors of log clade richness across time slices and trees (Fig. S7a, b; largest shared effects using standardized data). The percent of DNA-sampled species per clade is not important for explaining richness differences in multivariate models (Fig. S7c), indicating that completed mammal trees are unbiased for this question. In multivariate analyses (top three predictors across 100 mammal trees), we find that crown age has unique effects across all time slices, but not different to those found in 3 sets of RCBD simulated phylogenies (Fig. 4a, top panel). While crown age can nominally explain differences in mammal richness, its effect is no larger than expected if tree-wide speciation rates were constant through time. In contrast, tip DR mean explains more variation in mammal richness than expected (and double that of crown age), especially from 5–30 Ma (Fig. 4a, center panel; non-overlapping 95% confidence intervals [CIs] between mammal trees and simulations). Similarly, tip DR skew has unique effects on mammal richness that increase for older time slices, especially at >50 Ma (Fig. 4a, bottom panel). Taken together, these results show that (i) recent speciation-rate variation in mammals is greater than expected from RCBD rate stochasticity alone; (ii) clade richness differences are better explained by recent speciation rates (tip DR mean and skew) than origin times (crown ages); and (iii) variation in both rates and ages is nonetheless important for explaining richness. Named clades show mostly similar results, but lack a way to generate null RCBD expectations (Fig. S8).

### Ecological effects on speciation rates

We analyzed the shared effects of standardized species’ ecological traits on tip DR values across 1,000 mammal trees (Fig. 5a, tip-level PGLS for 5,675 extant non-marine species). We find that vagility is inversely related to tip DR such that lower vagility mammals have faster recent speciation rates, both overall and especially for herbivores (*N* = 1,637) and carnivores (*N* = 1,565) versus omnivores (*N* = 1,852). For diurnality, daytime-active mammal species (*N* = 1,037) also have faster recent speciation, especially for diurnal herbivores (*N* = 450) as compared to omnivores (*N* = 478) or carnivores (*N* = 109). Lastly, species’ absolute latitudinal centroids are unrelated to their tip DR values, either across all mammals or within trophic categories (*P* > 0.05 for nearly all comparisons; Fig. 5a). Sensitivity analyses show that these tip-level PGLS results are robust to a range of alternatives (Fig. S9): using tip-dated mammal trees, using node density for recent speciation rates, removing taxonomically imputed species from completed trees, and removing island endemic species.

**Fig. 5.**
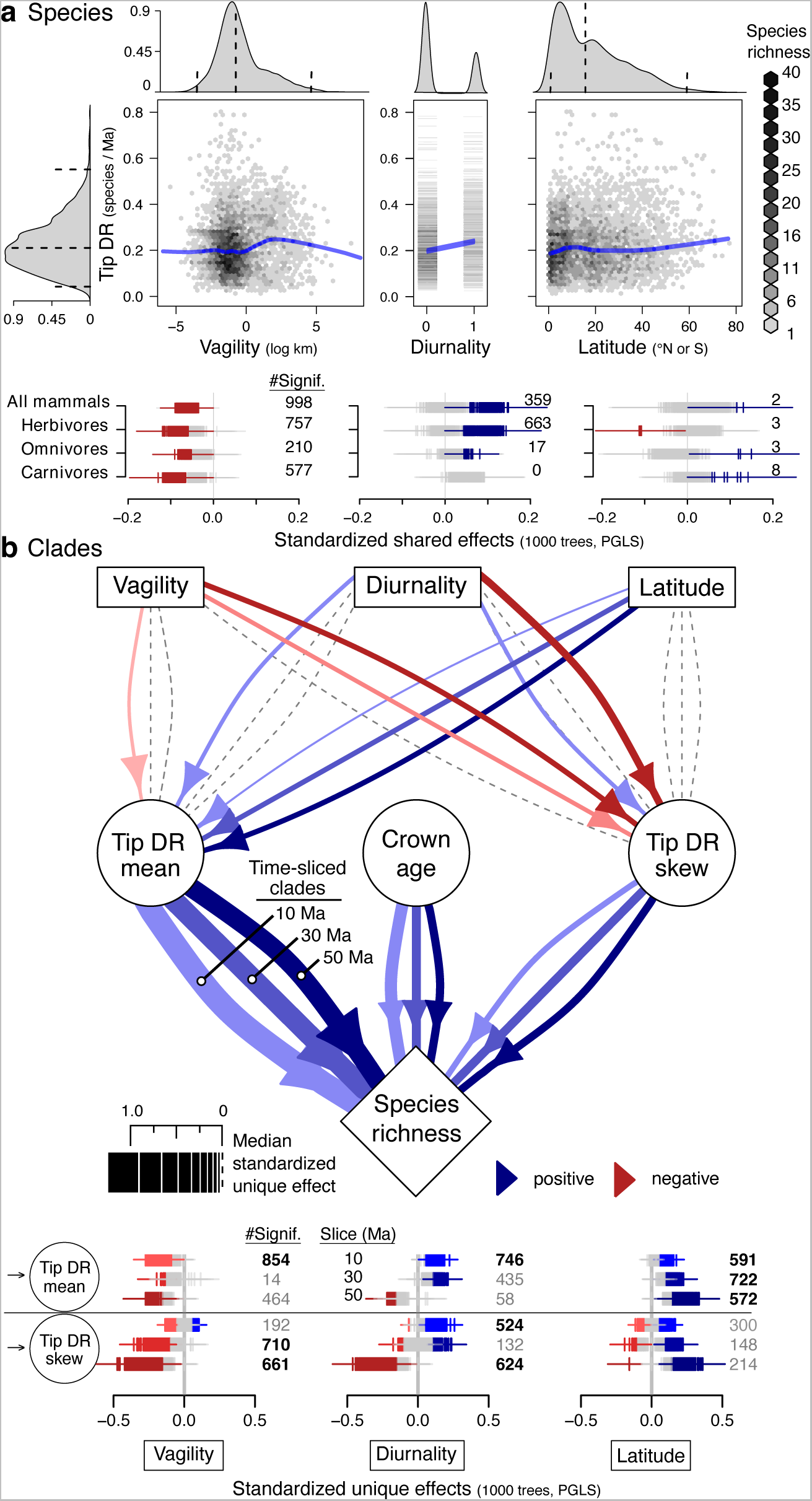
Connecting ecological and temporal causes of rate and richness variation in the mammal timetree. (**a**, top panels) Distribution of tip-level speciation rates (tip DR metric, shown is the harmonic mean of 10,000 trees) relative to per-species estimates of vagility (allometric index of maximum natal dispersal distance), diurnality (0=nocturnal or cathemeral, 1=diurnal), and absolute value of latitude (centroid of expert maps). Loess smoothing lines visualize general trends without considering phylogeny (blue, span=0.33). (**a**, bottom panels) Species-level effects considering phylogeny between tip DR and ecological attributes, as subset across trophic levels of herbivores, omnivores, and carnivores (univariate PGLS [phylogenetic generalized least squares] conducted on standardized predictors across 1,000 trees, showing 95% confidence intervals of slopes; colored if effects are significant, red for negative, blue for positive, else gray). (**b**) Phylogenetic path analysis conducted across time-sliced clades at 10-, 30-, and 50-Ma intervals, delimited as illustrated in Fig. 2. Path thickness, color, and directionality denote median coefficients of model-averaged analyses across 1,000 trees (time-sliced clades of 10-, 30-, and 50-Ma proceed from left to right as labeled). The bottom panels provide per-estimate uncertainty across time slices (slope ± SE), with non-zero estimates totaled as ‘#Signif.’ in the right margin. Paths present in >500 trees are bolded and displayed in the upper path model diagram whereas other paths are dashed lines.

Phylogenetic path analyses of time-sliced clades and including species attributes confirm that the unique effects of tip DR mean on clade richness are consistently ∼2x greater than crown age or tip DR skew (Fig. 5b), as found without considering attributes (Fig. 4a). We find that among 10-Ma clades, low vagility leads to faster rates of recent speciation (Fig. 5b), similar to the pattern seen at tree tips (Fig. 5a). Vagility effects are weaker among older clades, where vagility instead shows negative effects on tip DR skew (Fig. 5b), which is similar to the shared effects pattern seen in univariate PGLS (Fig. S10a). Diurnality shows positive effects on tip DR mean among 10-Ma clades, which is also analogous to the pattern at tree tips (Fig. 5a, b), and inconsistent effects on tip DR skew that are similar to the shared effects pattern (Fig. 5b; S10a). Latitude shows consistent, positive effects on tip DR mean among 10-Ma and older clades (Fig. 5b), which contrasts with the lack of tip-level latitudinal effect (Fig. 5a, S9) and inconsistent clade-level effects seen in univariate analyses (Fig. S10a). Both the time-sliced and taxon-based path analyses (Fig. S11) differ markedly from the univariate ecological analyses (Fig. S10), highlighting the importance of connecting both ecological and temporal factors in the investigation of uneven clade richness patterns. Sensitivity tests show that path analyses of time-sliced clades are robust to the exclusion of island endemics and imputed species, use of the tip-instead of node-dated backbone trees, and the use of node density instead of tip DR to summarize clade-level rate mean and skew (Fig. S12).

## DISCUSSION

Our investigations establish a primary role for ecological over temporal factors in causing uneven patterns of species richness in living mammals, suggesting that similar processes are likely also active in other branches of life. To reach this conclusion, we tested a hierarchical set of causal hypotheses (Fig. 1) for which types and speeds of macroevolutionary processes are occurring across phylogenetic levels, using an innovative time-slicing approach to define clades (Fig. 2). This framework enabled us to query evolutionary questions from shallow to deep levels of the extant timetree, recording macroevolutionary signals in recently diverged clades where inferences are more accurate and in more ancient clades where unobserved extinctions have accumulated (Kubo & Iwasa 1995; Marshall 2017; Louca & Pennell 2020; Upham *et al*. 2021). By connecting clade-level variation in ages and rates to species’ ecological attributes—both intrinsic (vagility, diurnality) and extrinsic (latitude)—this framework also connects two ideas that are usually investigated separately: phylogenetic unevenness (PU) and the latitudinal diversity gradient (LDG). Thus, applying this causal time-slice framework quantifies the relative roles of the PU, LDG, and intrinsic trait-rate processes in producing extant mammal biodiversity.

Considering varying clade ages, rates, and attributes of mammals (Fig. 3), we found evidence that speciation rates explain more of the variation in among-clade species richness than do crown ages (Fig. 4). This result refutes the idea that ‘clocklike’ rates of speciation predominate (contra (Ricklefs 2003; Venditti *et al*. 2010; Hedges *et al*. 2015)). We then ask why some clades have faster rates of speciation than others, finding that vagility and diurnality are greater causes of recent speciation-rate variation than is latitude (Fig. 5). Low-vagility and daytime-active species of mammals show the fastest recent speciation rates (supports H_A1_ and H_A2_, respectively), which suggests respective roles for dispersal limitation leading to peripatric speciation (Jablonski 1986; Kisel & Barraclough 2010) and diurnal adaptations leading to ecological speciation via time partitioning (Gerkema *et al*. 2013; Maor *et al*. 2017). In contrast, latitude positively affects speciation rates as measured in older clades, suggesting that faster speciation in temperate clades—coupled with the extinction of many nascent lineages—has helped produce the LDG pattern of greater tropical than temperate richness (supports H_A4_ over H_A3_). Thus, we here present evidence for contrasting modes of nonadaptive and adaptive speciation occurring in the same large radiation of mammals (*sensu* Czekanski-Moir & Rundell (2019)). We present arguments to justify these interpretations in the sections below.

### Time-sliced clades to test age and rate effects on richness

Original claims that uneven trees are random outcomes of constant-rate diversification (Wright 1941) have been refuted by several authors (Blum & François 2006; Davies *et al*. 2011; Rabosky *et al*. 2012), but with others continuing to support constant rates of speciation, extinction, or both (Ricklefs 2003; Venditti *et al*. 2010; Hedges *et al*. 2015). Ranked taxa have been the typical units for testing the relative effects of ages and rates on species richness differences. However, high variance in the crown ages of same-rank taxa (e.g., Table S1) has yielded proposals for time-standardizing higher taxa to make them comparably aged units (e.g., (Avise & Johns 1999; Dubois *et al*. 2021)). Time-standardization is both impractical due to regular phylogenetic flux (e.g., see (Franz *et al*. 2016, 2019)) and would disrupt the core principle of prevailing usage in taxonomic classification (Simpson 1945). Thus, we here embrace an alternative strategy of using tree-wide time slices to delimit comparable units of phylogenetic analysis (Fig. 2). A key feature of time-sliced clades is that they are readily delimited across a *sample of trees*—rather than a single consensus tree—so that clade sets on each tree can be iteratively used in comparative analyses. Doing so propagates confidence in node ages and relationships into modeled predictor effects, avoiding problems from assuming that the analyzed tree is the true tree (Huelsenbeck *et al*. 2000).

Under this time-slicing approach, we find a greater role for clade-level speciation rates (tip DR mean and skew) than time since most recent common ancestor (crown age) as the direct cause of mammal richness unevenness (Fig. 4a; Table 1). However, this explanatory role for speciation-rate variation is also greater than expected if Mammalia-wide birth and death rates were constant through time and among clades. Hence, these findings support arguments that ‘ecology’ (broadly defined as including any non-temporal factor that alters macroevolutionary-rate processes, including sexual selection and geographic factors) is a greater cause of species richness variation than is ‘time’ (here as crown age; (Price *et al*. 2012; Castro-Insua *et al*. 2018; Machac *et al*. 2018)). However, the ‘ecology’ vs. ‘time’ dichotomy is misleading. Variation in both rate and age clearly contribute to richness.

**Table 1.**
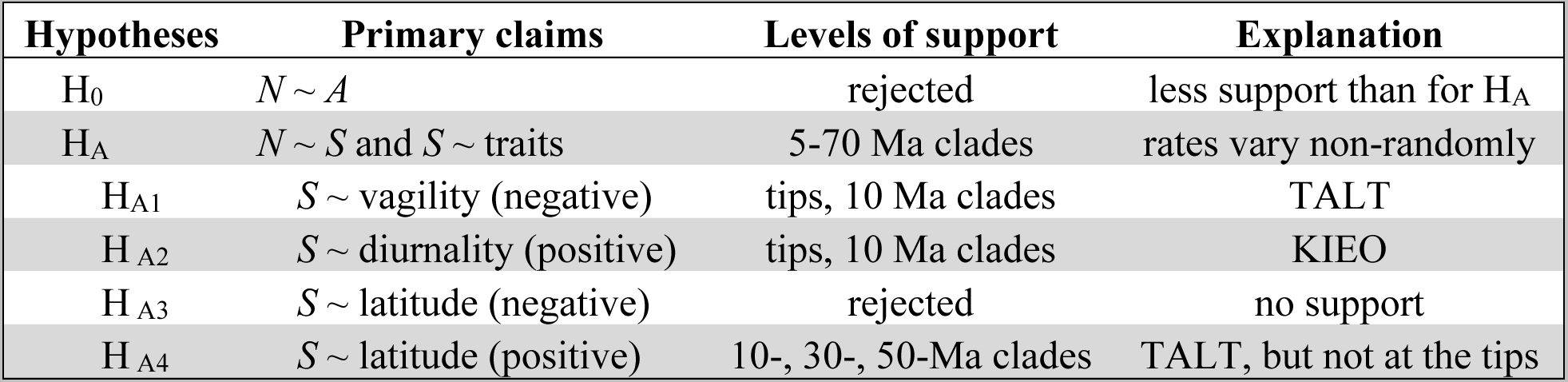
Summary of findings. Hypotheses refer to those in Fig. 1, and primary claims regarding how uneven clade richness (*N*) is explained by crown ages (*A*), speciation rates (*S*), and/or species’ attributes (traits). Levels of support refer to findings in Fig. 4 (H_0_ vs. H_A_) and Fig. 5 (H_A1-4_), while explanations include the interpretations of *trait-associated lineage turnover* (TALT) and *key innovation / ecological opportunity* (KIEO) discussed in the main text.

The dual roles of rate and age variation are intuitively illustrated by separating out those 10-Ma clades within 1 tree (Fig. 4b) and comparing subtrees of the most speciose clades (Fig. 4c). A rapid proliferation of species-level branches is needed for a given clade to be among the most speciose of a particular time slice, but some speciose clades are also near the 10-Ma maximum crown age for that example. Thus, while speciation-rate variation appears to cause the majority of the mammal PU pattern, it is only warranted to claim a greater *relative role* for ‘ecology’ over ‘time’ to the extent that rate-covarying ecological factors can also be identified.

### How do ecological differences lead to uneven speciation rates and richness?

Given the known greater influence of unsampled extinction events at deeper levels (older time slices) of the mammal timetree (Upham *et al*. 2021), we wanted to test how trait-to-speciation rate relationships differ depending on phylogenetic depth. We find that tip-level rate relationships with vagility (negative, H_A1_) and diurnality (positive, H_A2_) are only recovered for species’ tip rates and shallow 10-Ma clade rates, and not among clades at 30- or 50-Ma time slices (Fig. 5). This finding can be explained in two main ways. Either (i) the trait-to-rate signatures exist in older clades, but are invisible in the timetree due to unsampled extinct lineages; or (ii) the trait originated more recently than the 30- or 50-Ma cutoff point, so that the trait-to-rate signatures do not exist among older clades (younger signatures are swamped out in old clades).

In the first explanation, fast trait-associated speciation at the level of tips and 10-Ma clades is linked to similarly rapid extinction (*trait-associated lineage turnover* model; TALT). This view extends from models of ephemeral speciation, in which incipient species are frequently forming and going extinct (Mayr 1963; Stanley 1985; Rosenblum *et al*. 2012). However, here TALT further posits that lineage turnover is non-random with respect to certain trait or attribute states. In the second explanation, fast speciation among tips and young clades is a recent phenomenon caused by a trait adaptation that influences macroevolutionary rates (*key innovation / ecological opportunity* model; KIEO). This view stems from adaptive radiation theory, in which trait innovations can release certain lineages from competition, opening new ecological space that both promotes speciation and reduces extinction risk (Schluter 2000; Yoder *et al*. 2010; Gillespie *et al*. 2020). These contrasting TALT and KIEO views both posit that speciation rates have been higher in lineages with certain trait states, but they differ in their assumptions about unmeasured extinction rates (higher in TALT, lower in KIEO) and potentially the timing of historical onset in the trait-to-rate signatures (more recent onset of KIEO would explain the lack of older clade relationships, but TALT could have begun at any time). This TALT-or-KIEO framework therefore provides criteria for speculating about how species’ attributes are most likely influencing tree-wide speciation rates (Table 1).

### Vagility and turnover

The negative effects of vagility on tree-wide speciation rates indicate that low-vagility mammals—here proxied by smaller average body size and home/geographic ranges—are speciating faster than more vagile, wide-ranging species. The extremes of vagility within mammals help illustrate this dynamic, from low-vagility subterranean rodents like *Ctenomys* Tuco-Tucos—for which 68 range-restricted species are currently recognized ((MDD 2023); see Fig. 3, asterisk in clade 18 is the largest rate shift detected in Upham et al. (2021))—to the single species of highly vagile Mountain Lion, *Puma concolor*, which ranges from southern Alaska, USA to Tierra del Fuego, Argentina. This observed dynamic is consistent with long-standing theory linking individual-level dispersal ability (i.e., vagility) to lineage-level dynamics of gene flow versus isolation among geographic populations (Mayr 1963; Jablonski 1986; Slatkin 1987; Bohonak 1999; Kisel & Barraclough 2010). In this theory, less frequent dispersal leads to less gene flow and more genetic isolation in peripheral populations, promoting peripatric speciation (as distinct from allopatric speciation given the latter’s emphasis on physical barriers; (Mayr 1954; Carson & Templeton 1984)). Negative vagility-to-speciation rate relationships have been shown using the hand-wing index in birds (Claramunt *et al*. 2012; Sheard *et al*. 2020) and larval ecology in reef fishes (Riginos *et al*. 2014) and bivalves (Jablonski 1986). However, in the TALT-or-KIEO framework, we suggest that low-vagility lineages are apt to also experience high rates of *extinction* and thus high turnover (i.e., TALT). This is because the small ranges of low-vagility incipient species should present greater stochastic risk (Jablonski 1986; Kisel & Barraclough 2010), leading to the effects of vagility being erased at deeper levels of extant timetrees. If correct, the long branch leading to crown *Ctenomys* in the timetree (∼13 Ma; (Upham *et al*. 2019)) should be populated by many unsampled extinct taxa, a supposition that paleontological studies in fact support (6 genera of stem ctenomyids are known; (Verzi *et al*. 2013; De Santi *et al*. 2021)). Thus, we interpret the short-lived effects of vagility on speciation rates—only recorded at levels of tip species and shallow clades before subsiding—as evidence for TALT, implying that many low-vagility lineages older than 10 Ma have already gone extinct.

How, then, to explain the high rates of speciation detected in *Pteropus* Flying Foxes, which are among the largest-bodied and most vagile of all bats? We suggest that the influence of vagility on mammal diversification might be non-linear as hypothesized in birds (e.g., humped (Mayr 1963) or sigmoidal (Claramunt *et al*. 2012)), in which case our results among shallow clades and tip species may only be capturing one side of the vagility-to-rate relationship. The 57 living species of *Pteropus* (MDD 2023) originate from second highest rate shift in Upham et al. (2021) (Fig. 3, asterisk in clade 12). Their long-distance dispersal ability (Oleksy *et al*. 2015) has enabled *Pteropus* to reach most Indo-Pacific islands, a pattern of diversification that best fits a founder-event model of speciation (Tsang *et al*. 2020) and highlights the additional role of landscape heterogeneity in determining the shape of vagility-to-speciation rate relationships. Wing morphology is known to contribute to bat vagility (Norberg & Rayner 1987), similar to the hand-wing index in birds (Sheard *et al*. 2020), but bat flying abilities are not explicitly modeled by our vagility index. Instead, we rely on allometric scaling relationships as a rough proxy of dispersal distances across all mammals (Sutherland *et al*. 2000; Whitmee & Orme 2013). No morphological trait has yet been identified as a Mammalia-wide vagility metric, but physiological scaling relationships may offer additional resolution (e.g., the maximal sustainable metabolic rate divided by the metabolic cost of transport; (Hillman *et al*. 2014)). Nonetheless, the vagility patterns described here are robust to multiple sensitivity tests (including the exclusion of island endemics; Fig. S9, S12), and thus convey reliable macroevolutionary signatures of mammalian space use relative to speciation rates.

### Diurnality and persistence

In contrast, the KIEO perspective of adaptive diversification following a trait innovation best explains the observed pattern of faster speciation among diurnal tip species and 10-Ma clades and not thereafter (Fig. 5). The multiple independent origins of daytime activity started ∼35 Ma after a ‘nocturnal bottleneck’ among K-Pg-surviving mammals (Gerkema *et al*. 2013; Maor *et al*. 2017). This pattern of diurnal-associated speciation has been described at broader phylogenetic scales across major extant lineages of tetrapods (family-level sampling for mammals (Anderson & Wiens 2017)), as well as for narrower radiations of diurnal primates (Magnuson-Ford & Otto 2012; Santini *et al*. 2015; Arbour & Santana 2017) and whales (Morlon *et al*. 2011), but not before at the species level for all mammals. Fossil evidence suggests that non-mammalian synapsids may also have evolved diurnality (Angielczyk & Schmitz 2014), but those lineages are extinct and thus unobserved in the molecular timetree.

The coordinated eco-physiological changes required to evolve diurnality (e.g., eye pigments and corneal size (Gerkema *et al*. 2013)) have presumably carried with them fitness benefits from access to novel resources in the daytime niche. In this context, we posit that evolving diurnality has led to differential lineage persistence (i.e., low rates of species turnover = low extinction / high or moderate speciation) relative to nocturnality because novel niche resources have presumably improved organismal fitness (Yoder *et al*. 2010; Gerkema *et al*. 2013). The KIEO model implies that persistence-driven speciation—i.e., speciation rates that appear high in extant timetrees mainly because extinction rates are reduced—underlies the diurnal rate signature, in contrast to the turnover-driven speciation which we suggest is associated with low-vagility lineages and high latitudes. Interestingly, the acquisition of diurnal behavior has likely evolved and persisted at least ten times in crown mammals (Maor *et al*. 2017), from diurnal primates and squirrels to elephant shrews. However, there appears to be no characteristic secondary axis of resource specialization that is common across these groups (e.g., diet or locomotor diversity); rather, allopatric/peripatric speciation—and persistence of those isolated diurnal lineages—is likely the secondary driver of diurnal diversity (e.g., (Zelditch *et al*. 2015)). Overall, we show that faster diurnal than nocturnal speciation is a consistent signature among recent lineages of mammals, and suggest that it is caused by greater persistence (lower turnover) of lineages due to ecological opportunity in new daytime niches.

### Latitude and uneven clade-level speciation

Latitude-related hypotheses of climatic stability accelerating local adaptation and low-latitude speciation (H_A3_; (Mittelbach *et al*. 2007; Etienne *et al*. 2019)) or climatic instability spurring isolation and high-latitude speciation (H_A4_; (Cutter & Gray 2016; Schluter & Pennell 2017)) align with the aforementioned KIEO or TALT models of speciation, respectively. However, tropical net diversification must have been greater than at temperate latitudes to produce the LDG. We find no consistent influence of latitude upon tip-level speciation rates (Fig. 5a, S9), contrary to evidence from marine fishes (Rabosky *et al*. 2018) and angiosperms (Igea & Tanentzap 2020; Sun *et al*. 2020) that supported negative latitude-to-tip speciation rate relationships. Instead, we find only a clade-level inverse gradient in mammals: positive latitudinal effects on speciation rates are increasingly strong among 10-, 30-, and 50-Ma clades (Fig. 5b, S12) as well as among taxonomic orders (Fig. S11b). However, those clade-level latitudinal patterns are apparent only in multivariate path analyses (Fig. S10). Thus, our results bring nuance to the extant timetree-based perspective on the mammal LDG by showing that (i) the negative latitude-to-rate pattern is mainly present at deeper timetree levels (older clades); (ii) the pattern may be an artifact of unobserved extinctions biasing speciation-rate estimates in older clades; and (iii) whether artifact or not, the pattern is hidden unless covariation in other ecological causes of uneven speciation rates is considered.

Our latitudinal results compare to similarly mixed findings from other studies of birds and mammals. In timetrees of extant birds, latitude-to-tip rate effects are also absent at the global level (Jetz *et al*. 2012; Rabosky *et al*. 2015), but New World suboscines show faster temperate than tropical speciation (Kennedy *et al*. 2014; Harvey *et al*. 2020). New World sister species of birds and mammals similarly show higher turnover rates at temperate than tropical latitudes using mitochondrial DNA clocks (Weir & Schluter 2007; Schluter & Pennell 2017), but reliance on molecular clocks is more questionable for mammals than birds given their greater breadth of generation times (Nabholz *et al*. 2009). Other studies have shown inconsistent results with a variety of methods, including: (i) higher mammal subspecies counts in harsher temperate environments ((Botero *et al*. 2014); but the opposite pattern in birds (Martin & Tewksbury 2008)); (ii) no latitude-to-rate effects at the genus level (Soria-Carrasco & Castresana 2012) using consensus ages from the Bininda-Emonds et al. (2007) supertree of mammals; and (iii) greater rates of temperate extinction and tropical speciation (Rolland *et al*. 2014) using a modified version of that same mammal supertree. Here, using credible sets of mammal timetrees from the supermatrix-based Bayesian study of Upham et al. (2019) we find faster speciation rates in temperate than tropical clades, but weak relationships among tip species and shallow clades where speciation rates are most confidently inferred (Louca & Pennell 2020; Upham *et al*. 2021).

We speculate that the extinction-filtered lens of extant mammal diversity retains true signals of faster temperate than tropical lineage turnover, but that greater paleo-to-neontological synthesis will be needed to corroborate this pattern. Our use of species’ geographic range centroid distance from the equator is likely not capturing dependencies on environmental stability for widespread species with postglacial range expansions (e.g., red deer (Doan *et al*. 2022)). However, the latitudinal ‘essence’ of a given clade should be summarized increasingly well at deeper timetree levels by averaging the values of more modern species. This may be why we found the strongest latitude-to-rate patterns among 50 Ma clades and taxonomic orders. The ‘clades only’ pattern of faster high-latitude speciation is also biologically feasible if (i) high-latitude glaciations during the last ∼4 Ma (Mudelsee & Raymo 2005; Clague *et al*. 2020) led to a pulse of mammal extinctions, and (ii) those extinctions were phylogenetically dispersed enough to erase only the tip-level portion of the latitude-to-speciation rate effect. Under that scenario, clade-level signatures of faster temperate speciation are possible as long as temperate lineages were not fully extirpated during climatic oscillations (e.g., persisting in glacial refugia (Hewitt 2000)). This scenario is supported by the North American fossil record, in which mammal richness and latitude are not strongly correlated until ∼4 Ma (Marcot *et al*. 2016), as well as fossil evidence that high-latitude extinctions steepened the LDG for bivalves and reptiles (Jablonski *et al*. 2016; Meseguer & Condamine 2020). Overall, we contend that the traditionally invoked ‘cradle’ (higher tropical speciation) and ‘museum’ (lower tropical extinction (Mittelbach *et al*. 2007)) should instead re-focus upon the *turnover ratio* of those processes, more similar to Stebbens’ (1974) original meaning, as was emphasized by Vasconcelos et al. (2022). Testing whether mammal lineages have ‘cycled’ faster (i.e., shorter durations) outside than inside the tropics is the key question to resolve.

## CONCLUSION

By taking a broad view on the evolutionary history of Mammalia, from recent species’ tips to ancient clade-level processes, we uncover commonalities in the ecological causes of uneven species diversification over geography as well as phylogeny. We provide evidence that ecological factors have had non-random influences on mammalian macroevolutionary rates: far from stochastic, speciation rates are likely higher in clades that contain more low-vagility, diurnal, and temperate-distributed species. Ecologically linked speciation rates have, in turn, driven the majority of the PU pattern of uneven among-clade species richness. The LDG is not separate from this PU pattern, but rather interconnected with it—both types of unevenness share underlying causes, both direct (macroevolutionary rates and ages) and indirect (factors affecting gene flow, isolation, and adaptation). Species’ ecological attributes appear to provide the most reliable predictor for whether a given lineage will tend to speciate, go extinct, or persist in the near future (i.e., ‘species selection’ *sensu* Jablonski (2008)). Thus, the extent to which we can reliably unite the ecological, taxonomic, and phylogenetic knowledge of all mammal species— over 40% of which has only been described since 1993 (Burgin *et al*. 2018; MDD 2023)—is likely to dictate our ability to determine which aspects of extinction risk are inherent to species versus external and human-caused (Pyron & Pennell 2022).

Overall, we hypothesize that two main macroevolutionary processes are at work in mammals. First, we identify rate signatures that are consistent with turnover-driven speciation (TALT model) at shallow levels of the timetree due to greater geographic isolation among low-vagility species. Second, we hypothesize that persistence-driven speciation (KIEO model) is occurring in diurnal lineages because of access to new daytime niches and release from nocturnal competitors. We speculate different explanations for the same speciational signatures due to the differing circumstantial evidence of vagility acting on space use and diurnality acting on resource use. However, the value of applying the TALT-or-KIEO framework to our extant timetree-based analyses lies in it generating plausible hypotheses for more direct future evaluation using fossil-based extinction rates and species attributes. Developing physiological or skeletal metrics of mammalian vagility (as opposed to the allometric index used here) along with cranial correlates of diurnal vision (e.g., (Angielczyk & Schmitz 2014)) will be critical for assessing whether the relative frequency of turnover- and persistence-driven speciation has changed from fossil to modern ecosystems. Connecting evolutionary levels from individuals and species to clades appears promising for explaining uneven diversification across the tree of life.

## Acknowledgments

We thank the Upham Lab of Phylogenetic Ecology for discussions that improved this work. We also thank members of the VertLife Terrestrial grant (A. Pyron, G. Thomas, R. Bowie, R. Guralnick, M. Koo, D. Wake, T. Colston, M. Moura, A. Ranipeta) for initiating these efforts. Special thanks to I. Quintero, M. Landis, D. Schluter, A. Mooers, D. Greenberg, S. Upham and E. Florsheim for valuable conceptual input, as well as B. Patterson, K. Rowe, J. Brown, T. Peterson, D. Field, T. Stewart, and three anonymous reviewers for comments on earlier drafts of this study. M. Duong, G. Amatulli, and J. Hart provided technical support. Artwork from phylopic.org and open-source fonts. **Funding:** This work was supported by NSF VertLife Terrestrial grant to W.J. and J.A.E. (DEB 1441737 and 1441634), NSF grant DBI-1262600 to W.J., and NIH grant 1R21AI164268-01 and Arizona State University start-up funds to N.S.U. **Competing interests:** None.

## REFERENCES

Alroy, J. (2019). Small mammals have big tails in the tropics. Glob. Ecol. Biogeogr., 0.

Álvarez-Carretero, S., Tamuri, A.U., Battini, M., Nascimento, F.F., Carlisle, E., Asher, R.J., et al. (2022). A species-level timeline of mammal evolution integrating phylogenomic data. Nature, 602, 263–267.

Anderson, S.R. & Wiens, J.J. (2017). Out of the dark: 350 million years of conservatism and evolution in diel activity patterns in vertebrates. Evolution, 71, 1944–1959.

Angielczyk, K.D. & Schmitz, L. (2014). Nocturnality in synapsids predates the origin of mammals by over 100 million years. Proc. R. Soc. B Biol. Sci., 281, 20141642.

Arbour, J.H. & Santana, S.E. (2017). A major shift in diversification rate helps explain macroevolutionary patterns in primate species diversity. Evolution, 71, 1600–1613.

Avise, J.C. & Johns, G.C. (1999). Proposal for a standardized temporal scheme of biological classification for extant species. Proc. Natl. Acad. Sci., 96, 7358–7363.

Beaulieu, J.M. & O’Meara, B.C. (2016). Detecting Hidden Diversification Shifts in Models of Trait-Dependent Speciation and Extinction. Syst. Biol., 65, 583–601.

Belliure, Sorci, Møller, & Clobert. (2000). Dispersal distances predict subspecies richness in birds. J. Evol. Biol., 13, 480–487.

van der Bijl, W. (2018). phylopath: Easy phylogenetic path analysis in R. PeerJ, 6, e4718.

Bininda-Emonds, O.R.P., Cardillo, M., Jones, K.E., MacPhee, R.D.E., Beck, R.M.D., Grenyer, R., et al. (2007). The delayed rise of present-day mammals. Nature, 446, 507–512.

Blum, M.G.B. & François, O. (2006). Which Random Processes Describe the Tree of Life? A Large-Scale Study of Phylogenetic Tree Imbalance. Syst. Biol., 55, 685–691.

Bohonak, A.J. (1999). Dispersal, Gene Flow, and Population Structure. Q. Rev. Biol., 74, 21–45.

Bortolussi, N., Durand, E., Blum, M. & François, O. (2006). apTreeshape: statistical analysis of phylogenetic tree shape. Bioinformatics, 22, 363–364.

Botero, C.A., Dor, R., McCain, C.M. & Safran, R.J. (2014). Environmental harshness is positively correlated with intraspecific divergence in mammals and birds. Mol. Ecol., 23, 259–268.

Burgin, C.J., Colella, J.P., Kahn, P.L. & Upham, N.S. (2018). How many species of mammals are there? J. Mammal., 99, 1–14.

Carson, H.L. & Templeton, A.R. (1984). Genetic Revolutions in Relation to Speciation Phenomena: The Founding of New Populations, 36.

Castro-Insua, A., Gómez-Rodríguez, C., Wiens, J.J. & Baselga, A. (2018). Climatic niche divergence drives patterns of diversification and richness among mammal families. Sci. Rep., 8, 8781.

Clague, J.J., Barendregt, R.W., Menounos, B., Roberts, N.J., Rabassa, J., Martinez, O., et al. (2020). Pliocene and Early Pleistocene glaciation and landscape evolution on the Patagonian Steppe, Santa Cruz province, Argentina. Quat. Sci. Rev., 227, 105992.

Claramunt, S., Derryberry, E.P., Remsen, J.V. & Brumfield, R.T. (2012). High dispersal ability inhibits speciation in a continental radiation of passerine birds. Proc. R. Soc. Lond. B Biol. Sci., 279, 1567–1574.

Cutter, A.D. & Gray, J.C. (2016). Ephemeral ecological speciation and the latitudinal biodiversity gradient. Evolution, 70, 2171–2185.

Czekanski-Moir, J.E. & Rundell, R.J. (2019). The Ecology of Nonecological Speciation and Nonadaptive Radiations. Trends Ecol. Evol., 0.

Davies, T.J., Allen, A.P., Borda-de-Água, L., Regetz, J. & Melián, C.J. (2011). Neutral Biodiversity Theory Can Explain the Imbalance of Phylogenetic Trees but Not the Tempo of Their Diversification. Evolution, 65, 1841–1850.

De Santi, N.A., Verzi, D.H., Olivares, A.I., Piñero, P., Álvarez, A. & Morgan, C.C. (2021). A new Pleistocene *Ctenomys* and divergence dating of the hyperdiverse South American rodent family Ctenomyidae. J. Syst. Palaeontol., 19, 377–392.

Diaz, L.F.H., Harmon, L.J., Sugawara, M.T.C., Miller, E.T. & Pennell, M.W. (2019). Macroevolutionary diversification rates show time dependency. Proc. Natl. Acad. Sci., 201818058.

Doan, K., Niedziałkowska, M., Stefaniak, K., Sykut, M., Jędrzejewska, B., Ratajczak-Skrzatek, U., et al. (2022). Phylogenetics and phylogeography of red deer mtDNA lineages during the last 50 000 years in Eurasia. Zool. J. Linn. Soc., 194, 431–456.

Donati, G.F.A., Parravicini, V., Leprieur, F., Hagen, O., Gaboriau, T., Heine, C., et al. (2019). A process-based model supports an association between dispersal and the prevalence of species traits in tropical reef fish assemblages. Ecography, 42, 2095–2106.

Dubois, A., Ohler, A. & Pyron, R.A. (2021). New concepts and methods for phylogenetic taxonomy and nomenclature in zoology, exemplified by a new ranked cladonomy of recent amphibians (Lissamphibia). Megataxa, 5, 1–738.

Etienne, R.S., Cabral, J.S., Hagen, O., Hartig, F., Hurlbert, A.H., Pellissier, L., et al. (2019). A Minimal Model for the Latitudinal Diversity Gradient Suggests a Dominant Role for Ecological Limits. Am. Nat., 194, E122–E133.

Fine, P.V.A. & Ree, R.H. (2006). Evidence for a Time-Integrated Species-Area Effect on the Latitudinal Gradient in Tree Diversity. Am. Nat., 168, 796–804.

Foley, N.M., Mason, V.C., Harris, A.J., Bredemeyer, K.R., Damas, J., Lewin, H.A., et al. (2023). A genomic timescale for placental mammal evolution. Science, 380, eabl8189.

Franz, N.M., Musher, L.J., Brown, J.W., Yu, S. & Ludäscher, B. (2019). Verbalizing phylogenomic conflict: Representation of node congruence across competing reconstructions of the neoavian explosion. PLOS Comput. Biol., 15, e1006493.

Franz, N.M., Pier, N.M., Reeder, D.M., Chen, M., Yu, S., Kianmajd, P., et al. (2016). Two Influential Primate Classifications Logically Aligned. Syst. Biol., 65, 561–582.

Freckleton, R.P., Phillimore, A.B. & Pagel, M. (2008). Relating Traits to Diversification: A Simple Test. Am. Nat., 172, 102–115.

Gerkema, M.P., Davies, W.I.L., Foster, R.G., Menaker, M. & Hut, R.A. (2013). The nocturnal bottleneck and the evolution of activity patterns in mammals. Proc R Soc B, 280, 20130508.

Gillespie, R.G., Bennett, G.M., De Meester, L., Feder, J.L., Fleischer, R.C., Harmon, L.J., et al. (2020). Comparing Adaptive Radiations Across Space, Time, and Taxa. J. Hered., 111, 1–20.

Gittleman, J. l. & Purvis, A. (1998). Body size and species–richness in carnivores and primates. Proc. R. Soc. Lond. B Biol. Sci., 265, 113–119.

von Hardenberg, A. & Gonzalez-Voyer, A. (2013). Disentangling Evolutionary Cause-Effect Relationships with Phylogenetic Confirmatory Path Analysis. Evolution, 67, 378–387.

Harvey, M.G., Bravo, G.A., Claramunt, S., Cuervo, A.M., Derryberry, G.E., Battilana, J., et al. (2020). The evolution of a tropical biodiversity hotspot. Science, 370, 1343–1348.

Harvey, M.G. & Rabosky, D.L. (2018). Continuous traits and speciation rates: Alternatives to state-dependent diversification models. Methods Ecol. Evol., 9, 984–993.

Harvey, M.G., Seeholzer, G.F., Smith, B.T., Rabosky, D.L., Cuervo, A.M. & Brumfield, R.T. (2017). Positive association between population genetic differentiation and speciation rates in New World birds. Proc. Natl. Acad. Sci., 114, 6328–6333.

Hedges, S.B., Marin, J., Suleski, M., Paymer, M. & Kumar, S. (2015). Tree of life reveals clock-like speciation and diversification. *Mol. Biol. Evol.*, msv037.

Hewitt, G. (2000). The genetic legacy of the Quaternary ice ages. Nature, 405, 907–913.

Hillman, S.S., Drewes, R.C., Hedrick, M.S. & Hancock, T.V. (2014). Physiological vagility and its relationship to dispersal and neutral genetic heterogeneity in vertebrates. J. Exp. Biol., 217, 3356–3364.

Ho, L.S.T. & Ané, C. (2014). A Linear-Time Algorithm for Gaussian and Non-Gaussian Trait Evolution Models. Syst. Biol., 63, 397–408.

Huelsenbeck, J.P., Rannala, B. & Masly, J.P. (2000). Accommodating Phylogenetic Uncertainty in Evolutionary Studies. Science, 288, 2349–2350.

Igea, J. & Tanentzap, A.J. (2020). Angiosperm speciation cools down in the tropics. Ecol. Lett., 23, 692–700.

Isaac, N.J.B., Jones, K.E., Gittleman, J.L. & Purvis, A. (2005). Correlates of Species Richness in Mammals: Body Size, Life History, and Ecology. Am. Nat., 165, 600–607.

Jablonski, D. (1986). Larval ecology and macroevolution in marine invertebrates. Bull. Mar. Sci., 39, 565–587.

Jablonski, D. (2008). Species Selection: Theory and Data. Annu. Rev. Ecol. Evol. Syst., 39, 501– 524.

Jablonski, D., Huang, S., Roy, K. & Valentine, J.W. (2016). Shaping the Latitudinal Diversity Gradient: New Perspectives from a Synthesis of Paleobiology and Biogeography. Am. Nat., 189, 1–12.

Jablonski, D., Roy, K. & Valentine, J.W. (2006). Out of the Tropics: Evolutionary dynamics of the latitudinal diversity gradient. Science, 314, 102–106.

Jansson, R., Rodríguez-Castañeda, G. & Harding, L.E. (2013). What Can Multiple Phylogenies Say About the Latitudinal Diversity Gradient? A New Look at the Tropical Conservatism, Out of the Tropics, and Diversification Rate Hypotheses. Evolution, 67, 1741–1755.

Jetz, W., Carbone, C., Fulford, J. & Brown, J.H. (2004). The scaling of animal space use. Science, 306, 266–268.

Jetz, W. & Fine, P.V.A. (2012). Global Gradients in Vertebrate Diversity Predicted by Historical Area-Productivity Dynamics and Contemporary Environment. PLOS Biol., 10, e1001292.

Jetz, W., Thomas, G.H., Joy, J.B., Hartmann, K. & Mooers, A.O. (2012). The global diversity of birds in space and time. Nature, 491, 444–448.

Jones, K.E., Bielby, J., Cardillo, M., Fritz, S.A., O’Dell, J., Orme, C.D.L., et al. (2009).PanTHERIA: a species-level database of life history, ecology, and geography of extant and recently extinct mammals. Ecology, 90, 2648–2648.

Kennedy, J.D., Wang, Z., Weir, J.T., Rahbek, C., Fjeldså, J. & Price, T.D. (2014). Into and out of the tropics: the generation of the latitudinal gradient among New World passerine birds. J. Biogeogr., 41, 1746–1757.

Kisel, Y. & Barraclough, T.G. (2010). Speciation Has a Spatial Scale That Depends on Levels of Gene Flow. Am. Nat., 175, 316–334.

Kubo, T. & Iwasa, Y. (1995). Inferring the Rates of Branching and Extinction from Molecular Phylogenies. Evolution, 49, 694–704.

Louca, S. & Pennell, M.W. (2020). Extant timetrees are consistent with a myriad of diversification histories. Nature, 1–4.

Machac, A. & Graham, C.H. (2017). Regional Diversity and Diversification in Mammals. Am. Nat., 189, E1–E13.

Machac, A., Graham, C.H. & Storch, D. (2018). Ecological controls of mammalian diversification vary with phylogenetic scale. Glob. Ecol. Biogeogr., 27, 32–46.

Magnuson-Ford, K. & Otto, S.P. (2012). Linking the Investigations of Character Evolution and Species Diversification. Am. Nat., 180, 225–245.

Maor, R., Dayan, T., Ferguson-Gow, H. & Jones, K.E. (2017). Temporal niche expansion in mammals from a nocturnal ancestor after dinosaur extinction. *Nat*. Ecol. Evol., 1, 1889.

Marcot, J.D., Fox, D.L. & Niebuhr, S.R. (2016). Late Cenozoic onset of the latitudinal diversity gradient of North American mammals. Proc. Natl. Acad. Sci., 113, 7189–7194.

Marshall, C.R. (2017). Five palaeobiological laws needed to understand the evolution of the living biota. *Nat*. Ecol. Evol., 1, 1–6.

Martin, P.R. & Tewksbury, J.J. (2008). Latitudinal Variation in Subspecific Diversification of Birds. Evolution, 62, 2775–2788.

Mayr, E. (1954). Change of genetic environment and evolution. In: Evolution as a Process (eds. Huxley, J., Hardy, A.C. & Ford, E.B.). Allen & Unwin, London, pp. 157–180.

Mayr, E. (1963). *Animal species and evolution*. Belknap, Cambridge, MA.

McPeek, M.A. & Brown, J.M. (2007). Clade age and not diversification rate explains species richness among animal taxa. Am. Nat., 169.

MDD. (2023). Mammal Diversity Database v1.11.

Meredith, R.W., Janečka, J.E., Gatesy, J., Ryder, O.A., Fisher, C.A., Teeling, E.C., et al. (2011). Impacts of the Cretaceous Terrestrial Revolution and KPg Extinction on Mammal Diversification. Science, 334, 521–524.

Meseguer, A. & Condamine, F.L. (2020). Ancient tropical extinctions at high latitudes contributed to the latitudinal diversity gradient. Evolution, evo.13967.

Mittelbach, G.G., Schemske, D.W., Cornell, H.V., Allen, A.P., Brown, J.M., Bush, M.B., et al. (2007). Evolution and the latitudinal diversity gradient: speciation, extinction and biogeography. Ecol. Lett., 10, 315–331.

Moen, D.S. & Morlon, H. (2014). Why does diversification slow down? Trends Ecol. Evol., 29, 190–197.

Molina-Venegas, R. (2020). What are “tippy” and “stemmy” phylogenies? Resolving a phylogenetic terminological tangle. J. Syst. Evol., 59.

Mooers, A.O. & Heard, S.B. (1997). Inferring Evolutionary Process from Phylogenetic Tree Shape. Q. Rev. Biol., 72, 31–54.

Morlon, H., Parsons, T.L. & Plotkin, J.B. (2011). Reconciling molecular phylogenies with the fossil record. Proc. Natl. Acad. Sci., 108, 16327–16332.

Mudelsee, M. & Raymo, M.E. (2005). Slow dynamics of the Northern Hemisphere glaciation. Paleoceanography, 20.

Nabholz, B., Glémin, S. & Galtier, N. (2009). The erratic mitochondrial clock: variations of mutation rate, not population size, affect mtDNA diversity across birds and mammals. BMC Evol. Biol., 9, 54.

Norberg, U.M.L. & Rayner, J. (1987). Ecological morphology and flight in bats (Mammalia; Chiroptera): wing adaptations, flight performance, foraging strategy and echolocation. Philos. Trans. R. Soc. Lond. B Biol. Sci., 316, 335–427.

Oleksy, R., Racey, P.A. & Jones, G. (2015). High-resolution GPS tracking reveals habitat selection and the potential for long-distance seed dispersal by Madagascan flying foxes Pteropus rufus. Glob. Ecol. Conserv., 3, 678–692.

Phillimore, A.B. & Price, T.D. (2008). Density-dependent cladogenesis in birds. PLoS Biol., 6, e71.

Pontarp, M., Bunnefeld, L., Cabral, J.S., Etienne, R.S., Fritz, S.A., Gillespie, R., et al. (2019).The Latitudinal Diversity Gradient: Novel Understanding through Mechanistic Eco-evolutionary Models. Trends Ecol. Evol., 34, 211–223.

Price, S.A., Hopkins, S.B., Smith, K.K. & Roth, V.L. (2012). Tempo of trophic evolution and its impact on mammalian diversification. Proc. Natl. Acad. Sci. USA, 109, 7008–7012.

Purvis, A., Fritz, S.A., Rodríguez, J., Harvey, P.H. & Grenyer, R. (2011). The shape of mammalian phylogeny: patterns, processes and scales. Philos. Trans. R. Soc. Lond. B Biol. Sci., 366, 2462–2477.

Pyron, R.A. & Pennell, M. (2022). Macroevolutionary perspectives on Anthropocene extinction. Biol. Conserv., 274, 109733.

Quintero, I. & Jetz, W. (2018). Global elevational diversity and diversification of birds. Nature.

Rabosky, D.L. (2009). Ecological limits and diversification rate: alternative paradigms to explain the variation in species richness among clades and regions. Ecol. Lett., 12, 735–743.

Rabosky, D.L. (2014). Automatic Detection of Key Innovations, Rate Shifts, and Diversity-Dependence on Phylogenetic Trees. PLOS ONE, 9, e89543.

Rabosky, D.L. (2015). No substitute for real data: A cautionary note on the use of phylogenies from birth–death polytomy resolvers for downstream comparative analyses. Evolution, 69, 3207–3216.

Rabosky, D.L., Chang, J., Title, P.O., Cowman, P.F., Sallan, L., Friedman, M., et al. (2018). An inverse latitudinal gradient in speciation rate for marine fishes. Nature, 1.

Rabosky, D.L., Slater, G.J. & Alfaro, M.E. (2012). Clade age and species richness are decoupled across the eukaryotic tree of life. PLoS Biol., 10, e1001381.

Rabosky, D.L., Title, P.O. & Huang, H. (2015). Minimal effects of latitude on present-day speciation rates in New World birds. Proc R Soc B, 282, 20142889.

Revell, L.J. (2012). phytools: an R package for phylogenetic comparative biology (and other things). Methods Ecol. Evol., 3, 217–223.

Ricklefs, R.E. (2003). Global diversification rates of passerine birds. Proc. R. Soc. Lond. B-Biol. Sci., 270, 2285–2291.

Riginos, C., Buckley, Y.M., Blomberg, S.P., Treml, E.A., Heard, A.E.S.B. & Bronstein, E.J.L. (2014). Dispersal Capacity Predicts Both Population Genetic Structure and Species Richness in Reef Fishes. Am. Nat., 184, 52–64.

Rohlf, F.J., Chang, W.S., Sokal, R.R. & Kim, J. (1990). Accuracy of Estimated Phylogenies: Effects of Tree Topology and Evolutionary Model. Evolution, 44, 1671–1684.

Rolland, J., Condamine, F.L., Jiguet, F. & Morlon, H. (2014). Faster Speciation and Reduced Extinction in the Tropics Contribute to the Mammalian Latitudinal Diversity Gradient. PLOS Biol., 12, e1001775.

Rosenblum, E.B., Sarver, B.A.J., Brown, J.W., Roches, S.D., Hardwick, K.M., Hether, T.D., et al. (2012). Goldilocks Meets Santa Rosalia: An Ephemeral Speciation Model Explains Patterns of Diversification Across Time Scales. Evol. Biol., 39, 255–261.

Sánchez-Reyes, L.L., Morlon, H. & Magallón, S. (2017). Uncovering Higher-Taxon Diversification Dynamics from Clade Age and Species-Richness Data. Syst. Biol., 66, 367–378.

Santini, L., Rojas, D. & Donati, G. (2015). Evolving through day and night: origin and diversification of activity pattern in modern primates. Behav. Ecol., 26, 789–796.

Schluter, D. (2000). The ecology of adaptive radiation. Oxford University Press, Oxford, UK.

Schluter, D. & Pennell, M.W. (2017). Speciation gradients and the distribution of biodiversity. Nature, 546, 48–55.

Scholl, J.P. & Wiens, J.J. (2016). Diversification rates and species richness across the Tree of Life. Proc. R. Soc. B Biol. Sci., 283, 20161334.

Sheard, C., Neate-Clegg, M.H.C., Alioravainen, N., Jones, S.E.I., Vincent, C., MacGregor, H.E.A., et al. (2020). Ecological drivers of global gradients in avian dispersal inferred from wing morphology. Nat. Commun., 11, 2463.

Silvestro, D., Castiglione, S., Mondanaro, A., Serio, C., Melchionna, M., Piras, P., et al. (2020). A 450 million years long latitudinal gradient in age-dependent extinction. Ecol. Lett., 23, 439–446.

Simpson, G.G. (1945). The principles of classification and a classification of mammals. Bull. Am. Mus. Nat. Hist., 85, xvi-350 p.

Slatkin, M. (1987). Gene Flow and the Geographic Structure of Natural Populations. Science, 236, 787–792.

Soria-Carrasco, V. & Castresana, J. (2012). Diversification rates and the latitudinal gradient of diversity in mammals. Proc. R. Soc. B Biol. Sci., 279, 4148–4155.

Stanley, S.M. (1985). Rates of evolution. Paleobiology, 11, 13–26.

Stebbins, G.L. (1974). Flowering plants: evolution above the species level. Belknap Press of Harvard University Press, Cambridge, Mass.

Steel, M. & Mooers, A. (2010). The expected length of pendant and interior edges of a Yule tree. Appl. Math. Lett., 23, 1315–1319.

Sun, M., Folk, R.A., Gitzendanner, M.A., Soltis, P.S., Chen, Z., Soltis, D.E., et al. (2020). Recent accelerated diversification in rosids occurred outside the tropics. Nat. Commun., 11, 3333.

Sutherland, G.D., Harestad, A.S., Price, K. & Lertzman, K. (2000). Scaling of Natal Dispersal Distances in Terrestrial Birds and Mammals. Conserv. Ecol., 4, art16.

Title, P.O. & Rabosky, D.L. (2019). Tip rates, phylogenies and diversification: What are we estimating, and how good are the estimates? Methods Ecol. Evol., 10, 821–834.

Tsang, S.M., Wiantoro, S., Veluz, M.J., Sugita, N., Nguyen, Y.-L., Simmons, N.B., et al. (2020). Dispersal out of Wallacea spurs diversification of Pteropus flying foxes, the world’s largest bats (Mammalia: Chiroptera). J. Biogeogr., 47, 527–537.

Upham, N.S., Esselstyn, J.A. & Jetz, W. (2019). Inferring the mammal tree: species-level sets of phylogenies for questions in ecology, evolution, and conservation. PLOS Biol.

Upham, N.S., Esselstyn, J.A. & Jetz, W. (2021). Molecules and fossils tell distinct yet complementary stories of mammal diversification. Curr. Biol., 31, 4195–4206.e3.

Vasconcelos, T., O’Meara, B.C. & Beaulieu, J.M. (2022). Retiring “Cradles” and “Museums” of Biodiversity. Am. Nat., 199, 194–205.

Venditti, C., Meade, A. & Pagel, M. (2010). Phylogenies reveal new interpretation of speciation and the Red Queen. Nature, 463, 349–352.

Verzi, D.H., Olivares, A.I. & Morgan, C.C. (2013). Phylogeny and Evolutionary Patterns of South American Octodontoid Rodents. Acta Palaeontol. Pol., 59, 757–769.

Wei, T. (2017). An introduction to corrplot package. Available at: https://cran.r-project.org/web/packages/corrplot/vignettes/corrplot-intro.html. Last accessed 11 May 2018.

Weir, J.T. & Schluter, D. (2007). The Latitudinal Gradient in Recent Speciation and Extinction Rates of Birds and Mammals. Science, 315, 1574–1576.

Whitmee, S. & Orme, C.D.L. (2013). Predicting dispersal distance in mammals: a trait-based approach. J. Anim. Ecol., 82, 211–221.

Wiens, J.J. (2011). The causes of species richness patterns across space, time, and clades and the role of “ecological limits”. Q. Rev. Biol., 86, 75–96.

Willis, J.C. (1922). Age and Area. Cambridge University Press, Cambridge.

Wilman, H., Belmaker, J., Simpson, J., de la Rosa, C., Rivadeneira, M.M. & Jetz, W. (2014). EltonTraits 1.0: Species-level foraging attributes of the world’s birds and mammals. Ecology, 95, 2027–2027.

Wright, S. (1941). The “Age and Area” Concept Extended. Ecology, 22, 345–347.

Yoder, J.B., Clancey, E., Des Roches, S., Eastman, J.M., Gentry, L., Godsoe, W., et al. (2010). Ecological opportunity and the origin of adaptive radiations. J. Evol. Biol., 23, 1581– 1596.

Zelditch, M.L., Li, J., Tran, L.A.P. & Swiderski, D.L. (2015). Relationships of diversity, disparity, and their evolutionary rates in squirrels (Sciuridae). Evolution, 69, 1284–1300.

