## Supplementary Information for "Ecological causes of uneven mammal diversity"

**This PDF file includes:**

Table of Contents
Supplementary Methods
Supplementary Results
Figs. S1 to S12
Tables S1 to S3
Captions for Dataset S1
References for SI reference citations

**Other supplementary materials for this manuscript include the following:**

Dataset S1

**Table of Contents** (click to navigate)

### Supplementary Methods

1. Data

*1.1 Mammalian trait data*

Our workflow for gathering trait data for this study (Fig. S1) involved (i) unifying multiple trait taxonomies to our phylogeny's master taxonomy of 5911 species; and (ii) interpolating home range area and vagility to the species level for mammals. We unified data sources via a two-step process of matching binomial names to the synonym list of Meyer et al. (2015), and then matching those translated names directly to the master taxonomy. Using *join* and *awk* in the bash shell, we joined four databases: EltonTraits v1.0 (Wilman *et al.* 2014), IUCN Global Mammal Assessment (IUCN 2016), PanTHERIA (Jones *et al.* 2009), and the island rule database of Faurby and Svenning ((2016); called "F&S"). Dataset S7 gives our full trait database.

We focused on ten ecological variables (circles and rectangles in Fig. S2): body mass (log kg), home range (log $km^2$), geographic range (log $km^2$), vagility (log km), insularity (0/1), trophic level (1/2/3), diurnality (0/1), marine or not (0/1), latitude (centroid, deg), longitude (centroid, deg). Mean body masses were assembled per species from F&S (5351 species) and EltonTraits (46 species, only those coded with certainty level "1" for direct observations). We note that Smith (2003) is the citation for ~70% of those body mass values, which in turn gathered species means from the primary literature.

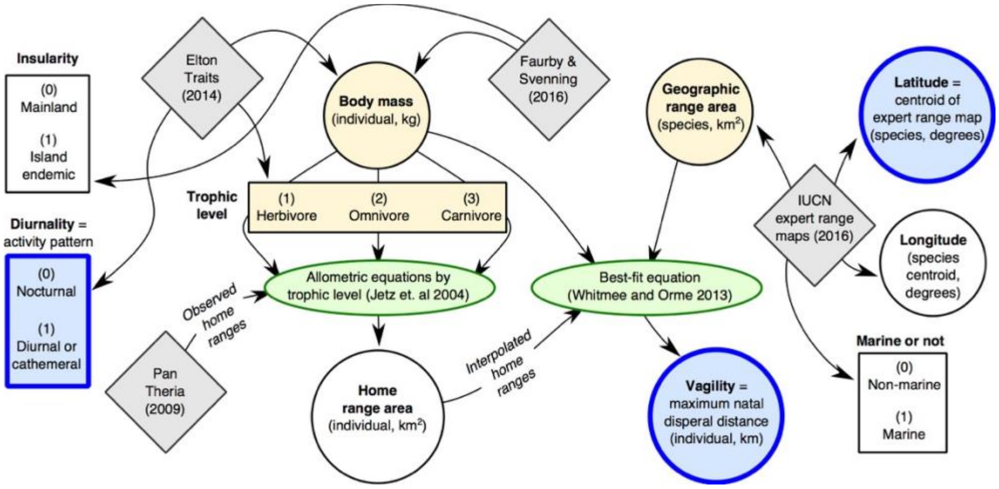

**Fig. S1**
**Workflow for trait data compilation.** (a) Schematic for compiling the 3 focal predictors (thick blue circles) of diversification-rate variation in main text Figs. 1 and 5. Shapes and colors represent: data sources (gray diamonds), interpolation equations (green ovals), continuous and discrete variables (circles and rectangles), and variables subjected to phylogenetic imputation (yellow fill). Coding categories are shown for discrete variables. Arrows represent input flows proceeding from data sources to taxonomic matching with our phylogeny.

The remaining 514 species without body mass values were imputed using the phylogeny and the R package "Rphylopars" (Goolsby *et al.* 2017). For each of 100 mammal trees, we (i) loaded the named trait data (including missing values); (ii) ran phylopars using models of Brownian motion (BM) and Pagel's lambda (LAM); and (iii) matched the model-imputed values back to the taxonomy. We then summarized the BM and LAM imputations across the 100 trees as medians and 95% CI bounds for each species. While the LAM models generally had better BIC scores than BM models, LAM yielded consistently larger values than BM, which were often unrealistic (e.g., *Acomys ngurui* has median of 573.3 grams [g] with LAM versus 90.9 g with BM, when the empirical range for *Acomys* is 18.5– 71.2 g; in contrast, the BM CI of 22.9–221.0 g is within the 13–38 g range given in the type description for *A. ngurui*, which was withheld (Verheyen *et al.* 2011)). We used the BM-based imputations of body mass for all calculations, including as related to interpolations of home range area and vagility (see below).

The geographic area and centroids of species geographic ranges were derived from expert range maps ((IUCN 2016), 18 August 2016 download). We calculated WGS84 ellipsoid areas of terrestrial and marine mammals in QGIS (QGIS Development Team 2017) while excluding the introduced portions of the ranges ("origin": 2, 3, or 4). Latitudinal and longitudinal centroids were taken as absolute values. Marine and non-marine mammals were given a binary coding. Insularity codings were gathered from F&S using the classical definition of island endemicity as pertaining to oceanic or landbridge islands (1122 species on islands of 5420 matched); species unmatched for island codings were manually interpolated (26 species on islands vs. 465 species assumed on the mainland). Missing values of geographic range area for 590 species were imputed based on phylogeny using the same Rphylopars procedure described above. Latitude and longitude centroid values were retained as "NA" for those 590 species without range maps due to concerns that geographic position is not a heritable trait in the same manner as body mass, geographic range, or intrinsic traits like activity pattern.

For trophic level, we modified the original diet classification from EltonTraits (10% intervals of 10 different food categories; (Wilman *et al.* 2014)) into categories of percent vertebrates, invertebrates, and plants. That re-categorization was then collapsed into an ordinal multistate variable per species: (1) herbivorous if 100% plant diet; (2) omnivorous if mix of plants and (in)vertebrates; and (3) carnivorous if 100% animal diet (vertebrates or not). Those 3 categories contained similar numbers of species (1637 vs. 1852 vs. 1565, respectively).

For activity pattern (diurnality), we similarly modified the overlapping binary codings of "nocturnal", "crepuscular", and "diurnal" in EltonTraits to make a mutually exclusive variable that was binary per species, as follows: (0) nocturnal if nocturnal only, nocturnal + crepuscular only, or crepuscular only; (1) diurnal if any daytime activity is present (i.e., diurnal only, diurnal + crepuscular, or mix of diurnal, crepuscular, and nocturnal). This re-categorization was clumped toward nocturnality (3413 vs. 1614 species, respectively). BM imputations of activity and trophic level were performed in Rphylopars the same as for continuous traits (imputing 857 species), except that median results were rounded to the nearest whole number to match missing species with values of their closest relatives.

Home range area per individual ($km^2$) was interpolated following the allometric equations of Jetz et al. (2004), which we updated using additional empirical data for non-marine home ranges reported in PanTheria ((Jones *et al.* 2009); n=603 species). Fig. S3 shows these empirical values best predicted by body masses (kg) with trophic level-specific equations ($R^2$ = 0.7–0.8 compared to 0.67 for all mammals). These equations can be compared directly to Table 1 in Jetz et al. (2004), although note that they reported slopes in $\log_{10}$ kg units while the intercepts are in non-log hectares (1 ha = 0.1 $km^2$) not the corresponding $\log_{10}$ $km^2$ units (see Fig. S3a). Our updated coefficients thus differ as follows: all mammals, 1.08 $\log_{10}$ kg slope (1.07 in Jetz et al. (2004)), 7.08 ha intercept (6.69); herbivores, 1.07 (1.02), 2.34 (2.05); omnivores, 1.14 (1.12), 12.02 (15.87); and carnivores, 1.35 (1.20), 29.51 (52.07). We used these updated trophic level equations to interpolate the home ranges of all remaining species (n=5308). We then validated model-predicted values vs. empirical observations (Fig. S3b).

Vagility, also called dispersal ability (e.g., (Claramunt *et al.* 2012)), was measured as the mean per-individual distance of maximum natal dispersal (km), and interpolated for each species following the best-fit equation of Whitmee and Orme (2013). That study identified the top 3 predictors of empirically measured maximum dispersal in mammals as adult body mass (g), home range size ($km^2$), and geographic range ($km^2$). They used natural log (ln) rather than $\log_{10}$, so we followed suit. We updated their equations using our estimates of the same traits by finding the best-fitting linear model to explain empirical data on maximum dispersal (n=89; (Whitmee & Orme 2013)). We estimated coefficients for the intercept, and slopes of body mass, home range, and geographic range as -2.496 (-1.153 in Whitmee and Orme (2013)), 0.206 (0.315), 0.323 (0.220), and 0.216 (0.252). We used that equation to interpolate estimates of maximum dispersal distance across all remaining species (n=5822), and validate predictions for the same 89 species as empirical data (Fig. S3b).

**a** Home range allometric equations

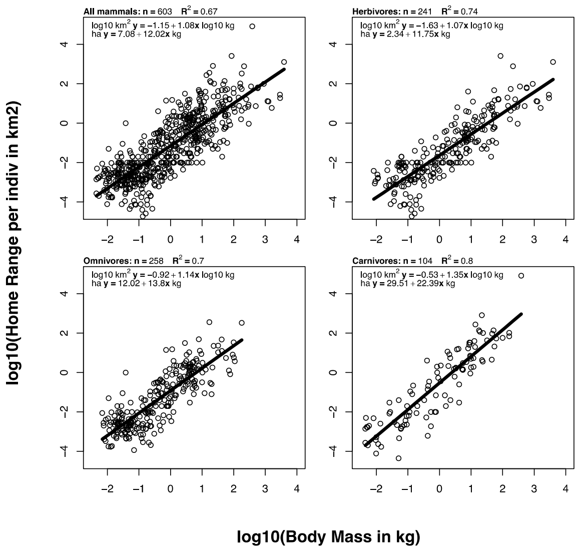

**b** Home range validation

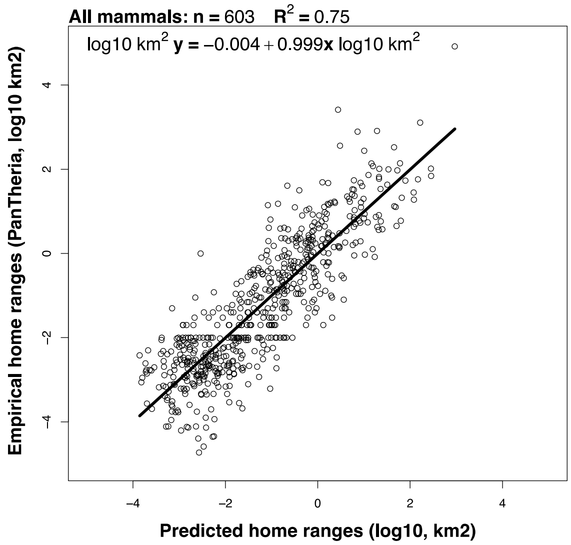

**c** Vagility index validation

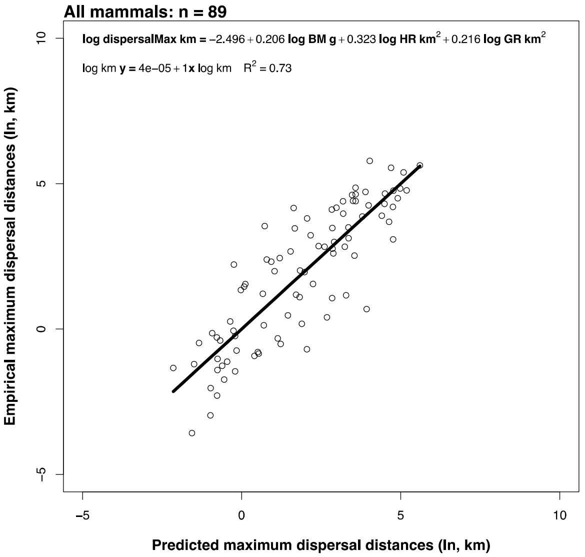

**Fig. S2**
**Data and equations used to predict per-individual estimates of home range and vagility.** (**a**) Using the approach of Jetz et al. (2004), we calculated log-log fits of body mass to empirically observed home range areas across all mammals (n=603) and within trophic-level categorizations. These updated per-trophic level equations were then used to expand the calculation of home range area to all species, which was in turn used in the vagility calculations (see Supplementary Information). (**b**) Shown is the resulting fit of our predicted data for those same empirical values (from the PanTheria database). (**c**) We used the top three predictors for maximum natal dispersal distance in the analysis of Whitmee and Orme (2013) to calculate log-log fits to updated empirical data. The resulting updated allometric equation was then used to expand the calculation of maximum dispersal distances to all species.

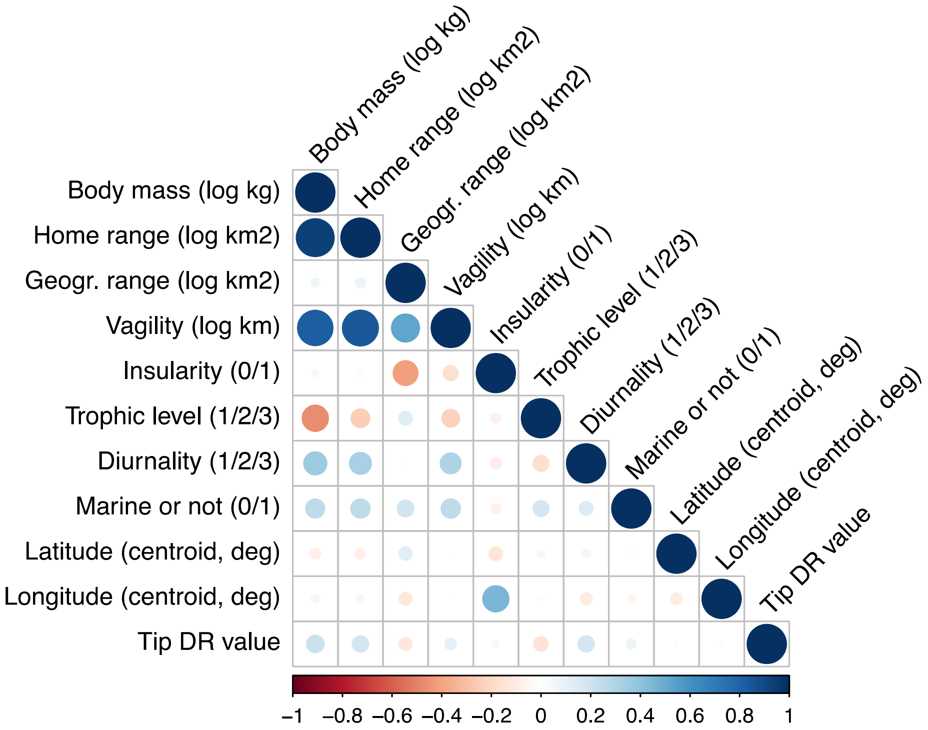

**Fig. S3**
**Trait data collinearity.** Pairwise spearman rank correlations among variables from which we selected independent predictors in analyses of speciation-rate variation. Colors correspond to correlation coefficients in the bottom legend. All extant species were used in these calculations (n=5,804). Tip DR values are here compared based on the harmonic mean per species of estimates from 10,000 completed trees. Note that these are non-phylogenetic correlations, so not interpretable in an evolutionary context.

#### *1.2 Tip-level diversification rates*

The primary means by which diversification rate variation was examined in this study was using per-species estimates of expected pure-birth (PB) diversification rates, calculated for the instantaneous present moment (tips of the tree) using the inverse of the equal splits measure (Redding & Mooers 2006; Jetz *et al.* 2012). We call this statistic "tip-level diversification rate" (tip DR) because it contains rate information weighted toward recent PB diversification processes occurring among extant species ((Quintero & Jetz 2018); we feel that "tip DR" is more descriptive than the term "DR statistic" used for this metric in Jetz et al (2012)). Redding and Mooers (2006) developed equal splits as a phylogenetic metric of per-lineage evolutionary isolation, Steel and Mooers (Steel & Mooers 2010) derived expected edge lengths in Yule trees, and Jetz et al (2012) applied the reciprocal of equal splits to measure recent rates of species PB diversification. The harmonic mean of tip DR closely approximates the PB diversification rate for clades greater than 10 species (supplementary equations 5 and 6 in Jetz et al (2012)). We calculate tip DR on full Mammalia phylogenies from the root to each tip as

$$Tip\ DR_i = 1 \Big/ \sum_{j=1}^{N_i} l_j \frac{1}{2^{j-1}}$$

where $N_i$ is the number of edges on the path from tip *i* to the root, and $l_j$ is the length of edge *j*. This equation assumes a fully bifurcating tree (Redding & Mooers 2006; Jetz *et al.* 2012). Because *j*=1 is the pendant edge leading to tip *i*, that branch length carries the greatest weight on the resulting value, with every ensuing rootward edge discounted exponentially as it is shared with other species. Sister species thus have identical tip DR values. Species with the highest tip DR have many short branches shared with other species toward the present, implying recent branching is abundant, whereas low-tip-DR species are subtended by long unshared branches (i.e., they are evolutionarily distinct (Redding & Mooers 2006)).

#### *1.3 Tip DR compared to model-based estimators*

Here we use tip DR in 2 ways: (i) at the species level, where tip DR measures recent rates of PB diversification (≈ speciation, if recent extinction is minimal); (ii) at the clade level, using the harmonic mean of tip DR among species (referred to as "tip DR mean") to approximate clade rates of PB diversification. The harmonic, not arithmetic, mean is preferred where rates are averaged (Ferger 1931). Fig. S1 shows that tip DR among species is closely related to tip net diversification rates from the BAMM birth-death model (see section 1.3, below). Tip speciation rates from BAMM are similarly highly correlated with species tip DR, while tip extinction rates are generally low (0–0.2 species/Ma) with a slight positive trend suggesting greater turnover in high-tip-DR species (Fig. S1a; see Quintero and Jetz (2018) for tip DR-to-BAMM comparisons in birds).

At the clade level, Fig. S1b shows that tip DR harmonic mean best approximates PB diversification rates across time-slice defined clades (see section 1.8). PB rates and tip DR mean show tighter correspondence for 50-Ma versus 10-Ma clades, as is expected since we calculated tip DR values upon full mammal trees but PB values on a per-clade basis (i.e., clades from older time slices are closer to the root, where tip DR mean equals the tree-wide Yule rate (Jetz *et al.* 2012)). Tip DR mean is thus a clade-level average of species-level branching processes that is meaningfully aligned with clade-level models of diversification rates that ignore extinction. As expected, tip DR mean underestimates birth-death (BD) rates of speciation and net diversification per clade (Fig. S1b), since BD processes are modelling extinct branches (largely unrecorded in our nearly extant-only mammal trees). Similar underestimates of BD rates by tip DR mean were found in named bird clades (supplementary methods Fig. 5d in Jetz et al (2012)) and bird elevational assemblages (fig. S6 in Quintero and Jetz (2018)). Additionally, we examined the skewness of clade tip DR (called "tip DR skew") as a way to characterize the clade tip DR distribution beyond its central tendency.

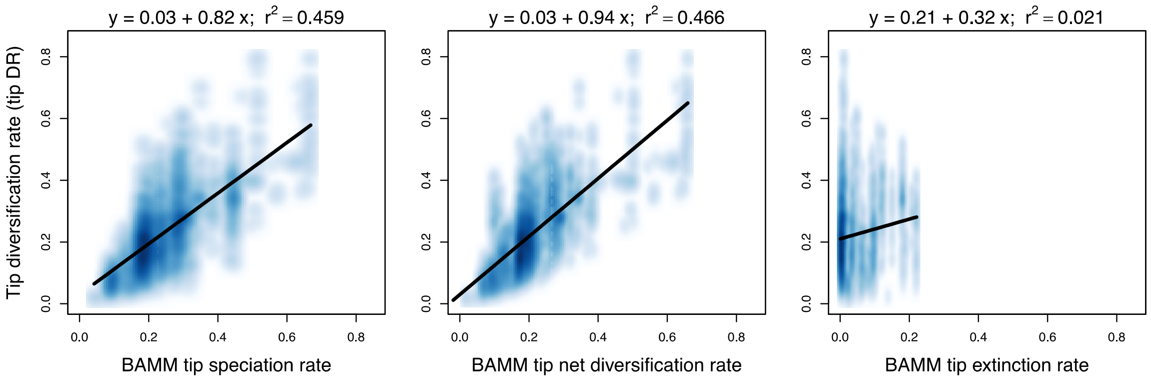

**Fig. S4**

**Comparison of tip-level diversification rates (tip DR) from this study to other rate metrics.** At the tip-level, species tip DR values are compared to tip-level estimates of speciation, net diversification, and extinction calculated in BAMM v2.5. Shown are the harmonic means of per-species values from 10,000 full Mammalia trees (node-dated backbone) versus BAMM tip rates (summary of runs on 10 trees). Note that BAMM tip rates for net diversification predict tip DR more strongly than does BAMM speciation rate (linear models shown).

### 2. Analyses

#### *2.1 Time-sliced clades*

In order to objectively define clades for tests of among clade variation in richness and diversification rates, we arbitrarily drew lines (referred to as “time slices”) at 5-Ma intervals and took the resulting *tipward* monophyletic clades as units of analysis. The *rootward* relationships of those clades (the “rootward backbone”) was retained for each interval, giving the expected covariance structure among clades when performing phylogenetic generalized least squares (PGLS) analyses. Fig. 3 gives a graphical example upon a subclade of the mammal tree. This procedure was scaled up to the full Mammalia tree—for example, for the 35-Ma slice, a typical tree resulted in 90 clades that varied in species richness from 2–726 species (mean, and standard error: 65.5, 13.0), and had crown ages from 0.4–34.9 Ma (22.5, 1.1). The key criterion uniting these crown clades is that their stem branch was extant at 35 Ma, so their species are living descendants of a lineage at least that age. We contrast this approach to analyses performed on named clades (see section 1.9).

Time-sliced clades were constructed at 14 slices from 5–70 Ma for each of 100 trees for Mammalia and the 3 sets of RC simulations (Fig. 3 shows the distribution of clade richness values across time slices). We used the “treeSlice” function in phytools with orientation set to “tipwards” and “rootwards” for time-slice clades and backbone, respectively, re-naming the backbone tips to match the row names of the tipward clade summaries. We summarized clades for the following values, with rates and ages specific to each tree: species richness; crown age (timing of first split not yet extinct, i.e., the most recent common ancestor [MRCA]); tip DR mean, skew, kurtosis, and coefficient of variation; PB and BD diversification rates, and BD speciation, extinction, and turnover; and percent of species sampled for DNA. By definition, all tipward clades at a time slice had identical stem ages. Singleton branches (“clades” of 1 species) were dropped from clade-level summaries, but retained on time-slice backbones with their species-level values of tip DR and richness for PGLS. Clades with 2 or more species were summarized for MRCA. Clades with four or more species were assessed for tip DR skew and PB and BD rates (n=4 is the minimum tree size for modelling rates).

Note that our use of the term “time slice” to delimit tipward clades and their rootward backbone differs from the usages of some other authors. Most commonly in palaeontology, only the rootward time slices of fossil chronograms are examined (e.g., (Harcourt-Brown 2002; Ruta *et al.* 2007; Bernardi *et al.* 2016)). That approach seeks to compare sets of contemporaneous lineages at past time slices, while considering present-day taxa representatives of the 0-Ma time slice (reviewed in (Tarver & Donoghue 2011)). In contrast, other authors have used time slices to delimit units of analysis *within* time intervals rather than tipward or rootward, summarizing rates or traits of tree-wide branches bounded by time bins (e.g., (Stadler 2011; Mazel *et al.* 2017)). Here we use time slices to delimit non-nested, tipward clades and their rootward phylogenetic covariance as an objective means of conducting clade-level PGLS analyses. We compare our results to analyses using traditional taxon-based clades (mammalian genera, families, and orders).

*2.2 Clade-level PGLS*

To test what factors best explain variation in species richness among clades, we performed PGLS analyses using Pagel's "lambda" transformation (Pagel 1994), as implemented in either the nlme R package (Pinheiro *et al.* 2018) or phylolm (Ho & Ané 2014). The phylolm package implemented the same PGLS models as nlme with numerically identical results in over an order of magnitude less time (results not shown), a finding that surprised us but is expected because phylolm is optimized for large trees (Ho & Ané 2014). As a result, we used phylolm wherever possible. We also used the caper package (Orme 2018) for PGLS when calculating $R^2$ values. Lambda was always estimated on a per tree and per comparison basis, so as to only correct for the amount of phylogenetic signal present in the data (lambda=1.0 is full Brownian motion). Extinct taxa were always excluded from PGLS.

We conducted PGLS upon time-slice clades for each of 100 trees as follows: (i) load in tree-specific backbones and tipward clade summaries; (ii) set the natural log of species richness as the response variable; (iii) standardize the predictors as mean centered and scaled by the standard deviation; and then for each slice interval (iv) match the backbone tree tips to the clade summary row names using "treedata" in geiger (Harmon *et al.* 2008); (v) perform univariate PGLS for each of the predictors to understand their shared effects on log richness; and (vi) select top predictors for multivariate PGLS to understand their unique effects on log richness. See Supplementary Results for results of the full univariate and multivariate analyses including percent sampling.

We repeated these 100-tree PGLS analyses for higher taxa to compare our results to other studies of clade richness in mammals (e.g., (McPeek & Brown 2007; Rabosky *et al.* 2012)). We used taxa having ≥ 4 species:

- Genera: n=385 of 1283 total, 4–192 species, 1.1–27.8 Ma crown age means across 100 trees (grand mean, standard error: 8.2, 0.3)
- Families: n=102 of 162 total, 4–768 species, 4.0–58.0 Ma (19.1, 1.2)
- Orders: n=22 of 27 total, 4–2354 species, 9.6–73.7 Ma (42.3, 4.4)

For the taxon-based clades, we similarly generated per-clade summaries of richness, MRCA, tip DR mean and skew, and PB and BD rates. Those summaries then served as data points united by the backbone structure of phylogenetic covariance between clades. Pruning clades off the backbone results is a non-ultrametric tree (due to their widely different stem ages), in contrast to the time-sliced clades where each slice produces a rootward ultrametric backbone. Our solution was to simply prune full Mammalia trees to per-clade representatives, and do it the same way for time-sliced and taxon-based clades, so that the pendant edges of clade backbones extended to the present day. Long terminal branch lengths do not affect the resulting phylogenetic covariance structure among clades, only internode relationships and distances do (Pagel 1994), so this procedure did not differently impact the PGLS analyses based on different clade delimitations. See Supplementary Results for univariate and multivariate predictions of species richness in taxa.

*2.3 Tip-level PGLS (correlates of diversification rates)*

For all PGLS analyses involving ecological traits (including those at the species level and clade level), we focused on a 5675-species data set that excluded (i) the 107 extinct species in our phylogeny due to trait uncertainty; and (ii) all 129 marine mammal species because of their order-of-magnitude trait differences relative to terrestrial species (and disparate selective constraints in marine environments; (Smith & Lyons 2013)). Sensitivity tests were additionally conducted for all rate-trait analyses to examine whether our conclusions were affected by the further exclusion of island endemic species (n=4553), DNA-imputed species (n=3941), and the separate analysis of trophic level categories (herbivore: n=1802; omnivore: n=2166; carnivore: n=1707 after excluding marine and extinct species; see Supplementary Results). For each of 100 trees, we used per-tree estimates of (natural) log tip DR, and species-level values for traits. We performed tip-level PGLS using the "phylolm" function in R, which performs well on large trees (Ho & Ané 2014), and plotted results using the "plotrix" (Lemon *et al.* 2017) and "hexbin" packages in R (Lewin-Koh 2018).

To better understand correlative structures underlying the observed rate variation across our mammal trees, we performed tip-level PGLS analyses between species' ecological traits and tip DR values across 100 trees. We followed Freckleton et al. (2008) in using trait ~ rate models in our tip-level PGLS analyses. This approach reverses the typical rate ~ trait model, where the rate is dependent on the independent trait (predictor) variable (Pagel 1997), even though the aim is still to draw inference on the effect of the trait upon the rate (Freckleton *et al.* 2008). This is because we sought to avoid identical residuals in the dependent variable — i.e., sister species by definition share all speciation events and have the same pendant branch length, so their tip DR values are always identical, which violates the assumption of within-variable data independence in bivariate normal distributions, and so should be avoided (Freckleton *et al.* 2008). The trait ~ rate approach was originally formulated for use with node depth (number of nodes from the root to each species' tip), but the same logic applies to other species-level rate metrics. Note that while sister species could share identical trait values, their independent evolution since splitting still arguably renders their traits independent (theoretically in this case, if not empirically).

Trait ~ rate PGLS has been applied to a variety of species-level evolutionary questions (e.g., (Matthews *et al.* 2011; Kozak & Wiens 2012; Dugo-Cota *et al.* 2015)), including with tip DR in univariate contexts (Harvey *et al.* 2017). Multivariate PGLS models (e.g., rate ~ trait1 + trait2) should not suffer the same issue, since the comparative method models the residual variation among trait variables in that case, not the rate ((Freckleton *et al.* 2008); see (Harvey *et al.* 2017) for an example using 9 traits). Although we did not conduct multivariate PGLS at the species level, we did do so at the clade level as part of phylogenetic path analyses upon time-sliced clades (see below, section 2.4)

*2.3 Clade-level PGLS (correlates of diversification rates)*

At the clade level, univariate PGLS could be performed typically, using rate ~ trait models, since averaging species' rates to clade tip DR mean resulted in sister clades having independent rate values. We performed univariate PGLS on time-sliced clades at 10-, 30-, and 50-Ma intervals as well as named taxonomic clades. These analyses were conducted by analogy to the previous analyses explaining log clade richness, with exceptions that (i) log tip DR mean was the dependent variable; and (ii) per-clade trait data summaries were the predictors (mean centered and standard deviation scaled to standardize the effect for comparison). Trait data was summarized using geometric means for vagility (to avoid skewing the clade means with large dispersal distances) and arithmetic means for latitudinal centroid (absolute value) and diurnality (averaging species coded 0–1 gave a clade proportion of diurnal species). Clade-level PGLS analyses were conducted on 100 trees (see Supplementary Results).

*2.4 Phylogenetic path analyses: clade-level causes of diversification and richness*

Path analyses are formal methods for translating between the languages of causality and statistical probability, the latter of which is usually employed in correlational (not causal) contexts. Causality can nonetheless be approached using statistics by completely resolving the correlational structure of a given phenomenon, which implies that all full and partial correlations underlying that phenomenon are known. Of the universe of theoretical correlation sets, only 1 can be causally correct. Sets are expressed as directed acyclic graphs (DAGs; also called structural equation models) and employed in path analyses to test the observed data (Shipley 2000).

Phylogenetic path analyses (PPA; (von Hardenberg & Gonzalez-Voyer 2013)) differ from other path models in their use of PGLS to test the statements of conditional independency that make up the "gaps" or non-paths in a path diagram (Gonzalez-Voyer & von Hardenberg 2014). For example, the model $A \rightarrow B \rightarrow C$ is made up of paths $C \sim B$, $B \sim A$, and $C \sim B + A$ (conditional dependencies), which means that several non-paths are also implied for this DAG to be true (conditional independencies). So if any of the non-paths (e.g., $C \sim A$) have significant PGLS slopes, it means that the null hypothesis of that independency is rejected, and thus the model $A \rightarrow B \rightarrow C$ is not the best fit to the observed data (path $A \rightarrow C$ may improve the model; (van der Bijl 2018)). Path analyses are generally confirmatory not exploratory (Shipley 2000), so a key step is constructing models to test the most relevant causal hypotheses.

We chose 27 path models for PPA using our observed mammal data, with the goal of connecting hypotheses of species' ecological traits impacting diversification rates and, in turn, patterns of observed clade richness (Fig. S5). All models had 3 levels of vertices consisting of (i) clade traits (log vagility, non-log diurnality and latitude; as described in section 2.3); (ii) clade rates (log tip DR mean and non-log tip DR skew) and log crown age; and (iii) log clade species richness. DAGs varied from having 0-3 paths from clade traits to rates, with each trait only able to affect 1 rate variable (mean or skew) at a time; no paths were drawn from clade traits to crown age (Fig. S5). All 27 models included the 3 paths from clade rates and ages to richness, as we previously showed is best supported (main text Fig. 4, Fig. S5). Time-sliced clades at 10-, 30-, and 50-Ma intervals were analyzed along with taxon-based clades. Sensitivity tests of path analyses were additionally conducted to examine the impact of excluding island endemic species (n=4553) and DNA-imputed species (n=3941; Supplementary Results).

We used the "phylopath" package in R (van der Bijl 2018) to specify the PPA models, as well as implement the PGLS analyses (conducted in phylolm) and compare models from the resulting outputs. For each of 1000 trees, we did the following: (i) load the tree and per-clade summaries for traits; (ii) subset to the targeted traits and standardize all model variables (including richness and rates) to establish common scaling; (iii) specify the 27 models; (iv) for each clade, prune the tree to the data using "treedata" in geiger (row names of clade summaries were given correspondences to tree tip labels in advance); and (v) use the "phylo_path" function to calculate model fits using PGLS and lambda transformations.

Our goal in summarizing results from the 27 models was to identify best supported *paths*, rather than 1 specific model over the others. For this reason, we used a model averaging approach focusing on AICc-equivalent scores called "CICc" to quantify model fits per tree and clade set (time-sliced or taxon-based clades). CICc uses Fisher's C-statistic to conduct goodness of fit tests on the conditional independencies of each model (Shipley 2013; Gonzalez-Voyer & von Hardenberg 2014). A given model passes the "d-sep test" if it has a C-statistic with $P > 0.05$ relative to the chi-square distribution with that degrees of freedom (Shipley 2009). However, for our time-slice analyses we found that only ~2% to 28% of models within 2 CICc units of the best model also satisfied the more conservative d-sep criterion (Supplementary Results). For this reason, we chose to focus only on the 2-CICc unit threshold for deciding which models would go to the next step of model averaging (had we used the d-sep criterion, not all trees and clade sets would have returned supported models). For each tree and clade set, we averaged path coefficients only if they appeared in a given model ("conditional" option in the phylopath function "average"; (von Hardenberg & Gonzalez-Voyer 2013; van der Bijl 2018)). We then took those 1000 averaged models and plotted each of their coefficients with standard error intervals, and took median coefficients of all 100 averages to plot in summary path diagrams (main text Fig. 5b, Supplementary Results).

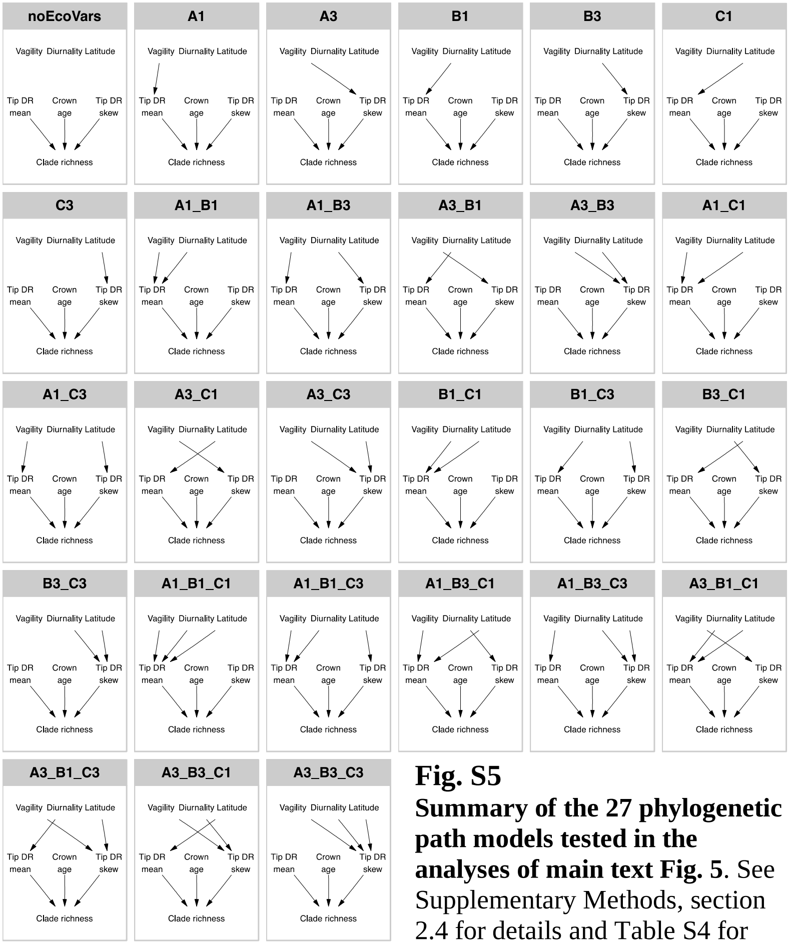

**Fig. S5**
**Summary of the 27 phylogenetic path models tested in the analyses of main text Fig. 5**. See Supplementary Methods, section 2.4 for details and Table S4 for modeling results.

### **Supplementary Results and Discussion**

#### 3. Supplementary Results

#### *3.1 Tip DR distribution across mammals*

We find fewer than expected low-tip-DR species (1st percentile) in the orders Rodentia ($\chi 2$ = 6.997, df = 1, P = 0.008), Primates ($\chi 2$ = 6. 308, df = 1, P = 0.012), and Artiodactyla ($\chi 2$ = 4.214, df = 1, P = 0.040), but no order contains more high-tip-DR species than expected (99th percentile; all P > 0.05).

#### *3.2 Characterizing time-sliced clades in mammals*

Tree-wide time slices at 10, 30, or 50 Ma produce a similar number of clades with comparable species richness as do mammal genera, families, or orders (Table S1). However, the crown ages of time-sliced clades vary ~50% less than do named clades in the same category, considering standard errors (Table 1). For example, mammal genera with ≥4 species have crown ages from 1.1–27.8 Ma across a sample of 100 phylogenies, while the comparable 10-Ma clades have crown ages from 0.2–9.9 Ma. Time-sliced clades are thus more time-uniform than named clades as a method of delimiting species diversity into comparable units for investigation.

**Table S1.**
**Comparison of clades with ≥4 species delimited by time slices versus taxonomic names.** Clades tipward of 10-, 30-, and 50-million year (Ma) intervals show less variation in crown age (smaller standard error, SE) than do named clades, as measured across 100 phylogenies from the credible set of node-dated mammal trees. See Fig. 2 main text for details of how time-sliced clades are delimited.

| | **Number of clades** | | **Species richness** | | **Crown ages (Ma)** | |
|---|---|---|---|---|---|---|
| **Clade** | **Mean** | **SE** | **Mean** | **SE** | **Mean** | **SE** |
| Time slices | | | | | | |
| 10 Ma | 387.9 | 1.6 | 12.9 | 0.7 | 7.5 | 0.1 |
| 30 Ma | 95.4 | 0.6 | 61.5 | 10.2 | 20.4 | 0.7 |
| 50 Ma | 39.3 | 0.5 | 152.1 | 43.6 | 34.5 | 2.0 |
| Taxa | | | | | | |
| Genera | 385 | N/A | 11.7 | 0.8 | 8.2 | 0.3 |
| Families | 102 | N/A | 56.1 | 12.0 | 19.1 | 1.2 |
| Orders | 22 | N/A | 263.5 | 117.1 | 42.3 | 4.4 |

Summarizing time-sliced clades at 5-Ma intervals from 5–70 Ma, it is clear that mammal phylogenies have a different shape than expected under homogeneous models of rate-constant birth and death (RCBD; Fig. S6). Mammal clades are more speciose than simulated clades from RCBD trees at every time slice examined (Fig. S6a), indicating that mammal trees are more "stemmy" with more extant tip-level diversity than expected (Rohlf *et al.* 1990; Molina-Venegas 2020). Extant species are also more heterogeneously distributed among clades than expected (greater richness variance; (Fig. S6b)), which is evidence for among-clade variation in crown ages, rates of speciation/extinction, or both.

#### *3.3 Comparing time-sliced and taxon-based clades in mammals*

By comparison to taxon-based clades, we find disparate patterns when conducting the same path analyses (Fig. S11), including: (i) no unique effects of vagility on tip DR mean in genera, families, or orders (versus inverse in 10-Ma clades); (ii) genera and families with negative vagility effects on tip DR skew (versus in 30- to 50-Ma clades); (iii) genera and families with positive diurnality effects on tip DR mean (versus only on 10-Ma clades); and (iv) orders show positive latitudinal effects on tip DR mean and skew (versus only on tip DR mean in time-sliced clades).

### 4. Sensitivity Tests

#### *4.1 Motivation to do sensitivity tests*

We conducted re-analyses of tip- and clade-level trait diversification on 1,000 trees from both the node- and tip-dated backbone distributions, in each case using data subsets to test for the following types of bias:

(i) *Excluding island endemic species* (n=4,553 non-marine species remaining): if islands have smaller geographic-range species (Brown *et al.* 1996) and stronger selective regimes on body size (Lomolino 2005), then we may expect them to bias our allometric calculations of vagility (Whitmee & Orme 2013), perhaps driving inverse vagility ~ tip DR in 10 Ma clades;

(ii) *Excluding DNA-missing species* (n=3,941 non-marine species remaining): if the birth-death polytomy resolver (PASTIS (Thomas *et al.* 2013)) that we used to impute DNA-missing species to our phylogeny creates a bias for trait evolution studies (Rabosky 2015), then excluding imputed species should change our tip- and clade-level results.

(iii) *Use of an alternative tip rates metric, node density (ND)*: if the reliance on tip DR is driving our results, then we expect key findings to change using another tip rate metric.

We were motivated to test for the influence of DNA-missing species imputed in trees on our analyses of trait diversification following the concerns raised by Rabosky (2015). Our comparison of results using ND (i.e., a simple count of nodes from the root to each species' tip (Freckleton *et al.* 2008)) was similarly motivated by Harvey and Rabosky (2018), who showed that PGLS using ND can in some cases have higher power than tip DR-based PGLS, though with a higher false discovery rate (0.06 vs. 0.02 doing PGLS with tip DR). Note that we did not conduct the 'sim' tests described in Harvey and Rabosky (2018) over concern that their use of rate ~ trait models violates the assumption of data point independence in bivariate normal distributions (sister species have identical tip rates (Freckleton *et al.* 2008); see Supplementary Methods section 2.3). Instead, we employed trait ~ rate PGLS models throughout our sensitivity tests that are nevertheless interpretable in the context of assessing trait-dependent diversification (see (Freckleton *et al.* 2008; Matthews *et al.* 2011; Harvey *et al.* 2017)). Overall, these sensitivity tests were broadly self-consistent and produced limited differences in the number of significant trees across backbone samples and data subsets.

*4.2 Tip-level PGLS*

The inverse effect of vagility ~ rate in 10-Ma clades is recovered similarly using the ND-based approach as using tip DR in both island and DNA-missing exclusions, as well as the trophic level subsets (Fig. S9, S12). Similarly, the mostly null tip-level effects of diurnality and latitude are consistent across subsets. The major difference is that the ND approach is less sensitive to the exclusion of DNA-imputed species – e.g., the 1,000 trees with significant vagility effects on ND for all mammals was reduced to 993 (and 992 tip-dated) versus being reduced from 998 to 60 (and 112) using tip DR. The ND approach thus usefully corroborates our tip DR-based findings and suggests that including the traits of DNA-missing species does not bias the conclusions of our tip DR analyses. DNA-missing species are in fact a non-random group of species – so excluding them may instead be a greater bias. Nevertheless, we agree that by testing for the sensitivity of our results to these imputed species we have confirmed the robustness of our trait diversification analyses.

*4.3 Clade-level path analyses*

Our results similarly hold at the level of time-sliced clades. Repeating these sensitivity analyses for path models finds qualitatively identical results, with minor differences in the number of trees for which a given path coefficient was non-zero (Fig. S11). For ease of interpretation, we summarize these data in Table S2.

For the inverse relationship of vagility ~ rate in 10-Ma clades, we found consistently strong evidence that excluding island and imputed species has no major influence (marginal relationships in the tip-dated backbone using tip DR mean; Table S3). Similarly, the positive diurnality ~ rate relationships in 10-Ma clades are consistently strong or marginal across nearly all comparisons. For latitude ~ rate, the tip DR-based comparisons reveal consistently strong or marginal results across the 10-, 30-, and 50-Ma clades, while the ND-based comparisons only have a consistent signal for the 50-Ma clades. All together, we highlight that the directionality and magnitude of results is visually consistent throughout (Fig. S11), an impressive finding given the extensive data and complex relationships explored. These sensitivity tests add rigor to our primary conclusions regarding ecological causes underlying the uneven recent diversification and species richness in mammals.

### Supplementary Data

**Data S1. (separate file)**
Species ecological trait data and code for analyses.

**Supplementary References**

Bernardi, M., Angielczyk, K.D., Mitchell, J.S. & Ruta, M. (2016). Phylogenetic Stability, Tree Shape, and Character Compatibility: A Case Study Using Early Tetrapods. *Syst. Biol.*, 65, 737–758.

van der Bijl, W. (2018). phylopath: Easy phylogenetic path analysis in R. *PeerJ*, 6, e4718.

Brown, J.H., Stevens, G.C. & Kaufman, D.M. (1996). The geographic range: Size, Shape, Boundaries, and Internal Structure. *Annu. Rev. Ecol. Syst.*, 27, 597–623.

Claramunt, S., Derryberry, E.P., Remsen, J.V. & Brumfield, R.T. (2012). High dispersal ability inhibits speciation in a continental radiation of passerine birds. *Proc. R. Soc. Lond. B Biol. Sci.*, 279, 1567–1574.

Dugo-Cota, Á., Castroviejo-Fisher, S., Vilà, C. & Gonzalez-Voyer, A. (2015). A test of the integrated evolutionary speed hypothesis in a Neotropical amphibian radiation. *Glob. Ecol. Biogeogr.*, 24, 804–813.

Faurby, S. & Svenning, J.-C. (2016). Resurrection of the Island Rule: Human-Driven Extinctions Have Obscured a Basic Evolutionary Pattern. *Am. Nat.*, 187, 812–820.

Ferger, W.F. (1931). The Nature and Use of the Harmonic Mean. *J. Am. Stat. Assoc.*, 26, 36–40.

Freckleton, R.P., Phillimore, A.B. & Pagel, M. (2008). Relating Traits to Diversification: A Simple Test. *Am. Nat.*, 172, 102–115.

Gonzalez-Voyer, A. & von Hardenberg, A. (2014). An Introduction to Phylogenetic Path Analysis. In: *Modern Phylogenetic Comparative Methods and Their Application in Evolutionary Biology* (ed. Garamszegi, L.Z.). Springer Berlin Heidelberg, Berlin, Heidelberg, pp. 201–229.

Goolsby, E.W., Bruggeman, J. & Ané, C. (2017). Rphylopars: fast multivariate phylogenetic comparative methods for missing data and within-species variation. *Methods Ecol. Evol.*, 8, 22–27.

Harcourt-Brown, K.G. (2002). Tree Balance, Time Slices, and Evolutionary Turnover in Cretaceous Planktonic Foraminifera. *Syst. Biol.*, 51, 908–916.

von Hardenberg, A. & Gonzalez-Voyer, A. (2013). Disentangling Evolutionary Cause-Effect Relationships with Phylogenetic Confirmatory Path Analysis. *Evolution*, 67, 378–387.

Harmon, L.J., Weir, J.T., Brock, C.D., Glor, R.E. & Challenger, W. (2008). GEIGER: investigating evolutionary radiations. *Bioinformatics*, 24, 129–131.

Harvey, M.G. & Rabosky, D.L. (2018). Continuous traits and speciation rates: Alternatives to state-dependent diversification models. *Methods Ecol. Evol.*, 9, 984–993.

Harvey, M.G., Seeholzer, G.F., Smith, B.T., Rabosky, D.L., Cuervo, A.M. & Brumfield, R.T. (2017). Positive association between population genetic differentiation and speciation rates in New World birds. *Proc. Natl. Acad. Sci.*, 114, 6328–6333.

Ho, L.S.T. & Ané, C. (2014). A Linear-Time Algorithm for Gaussian and Non-Gaussian Trait Evolution Models. *Syst. Biol.*, 63, 397–408.

IUCN. (2016). *IUCN RedList of Threatened Species. Version 2016.2*. Available at: http://www.iucnredlist.org/initiatives/mammals. Last accessed .

Jetz, W., Carbone, C., Fulford, J. & Brown, J.H. (2004). The scaling of animal space use. *Science*, 306, 266–268.

Jetz, W., Thomas, G.H., Joy, J.B., Hartmann, K. & Mooers, A.O. (2012). The global diversity of birds in space and time. *Nature*, 491, 444–448.

Jones, K.E., Bielby, J., Cardillo, M., Fritz, S.A., O'Dell, J., Orme, C.D.L., *et al.* (2009). PanTHERIA: a species-level database of life history, ecology, and geography of extant and recently extinct mammals. *Ecology*, 90, 2648–2648.

Kozak, K.H. & Wiens, J.J. (2012). Phylogeny, ecology, and the origins of climate–richness relationships. *Ecology*, 93, S167–S181.

Lemon, J., Bolker, B. & Oom, S. (2017). *Package "plotrix."* Available at: https://cran.r-project.org/web/packages/plotrix/plotrix.pdf. Last accessed 11 May 2018.

Lewin-Koh, N. (2018). *Hexagon binning: an overview*. Available at: https://cran.r-project.org/web/packages/hexbin/vignettes/hexagon_binning.pdf. Last accessed 11 May 2018.

Lomolino, M.V. (2005). Body size evolution in insular vertebrates: generality of the island rule. *J. Biogeogr.*, 32, 1683–1699.

Matthews, L.J., Arnold, C., Machanda, Z. & Nunn, C.L. (2011). Primate extinction risk and historical patterns of speciation and extinction in relation to body mass. *Proc. R. Soc. Lond. B Biol. Sci.*, 278, 1256–1263.

Mazel, F., Wüest, R.O., Lessard, J.-P., Renaud, J., Ficetola, G.F., Lavergne, S., *et al.* (2017). Global patterns of β-diversity along the phylogenetic time-scale: The role of climate and plate tectonics. *Glob. Ecol. Biogeogr.*, 26, 1211–1221.

McPeek, M.A. & Brown, J.M. (2007). Clade age and not diversification rate explains species richness among animal taxa. *Am. Nat.*, 169.

Meyer, C., Kreft, H., Guralnick, R. & Jetz, W. (2015). Global priorities for an effective information basis of biodiversity distributions. *Nat. Commun.*, 6, 8221.

Molina-Venegas, R. (2020). What are "tippy" and "stemmy" phylogenies? Resolving a phylogenetic terminological tangle. *J. Syst. Evol.*, 59.

Orme, C.D.L. (2018). The Caper Package: Comparative Analysis of Phylogenetics and Evolution in R.

Pagel, M. (1994). Detecting correlated evolution on phylogenies: a general method for the comparative analysis of discrete characters. *Proc R Soc Lond B*, 255, 37–45.

Pagel, M. (1997). Inferring evolutionary processes from phylogenies. *Zool. Scr.*, 26, 331–348.

Pinheiro, J., Bates, D., DebRoy, S., Sarkar, D. & R Core Team. (2018). nlme: Linear and Nonlinear Mixed Effects Models.

QGIS Development Team. (2017). QGIS Geographic Information System.

Quintero, I. & Jetz, W. (2018). Global elevational diversity and diversification of birds. *Nature*.

Rabosky, D.L. (2015). No substitute for real data: A cautionary note on the use of phylogenies from birth–death polytomy resolvers for downstream comparative analyses. *Evolution*, 69, 3207–3216.

Rabosky, D.L., Slater, G.J. & Alfaro, M.E. (2012). Clade age and species richness are decoupled across the eukaryotic tree of life. *PLoS Biol.*, 10, e1001381.

Redding, D.W. & Mooers, A.Ø. (2006). Incorporating Evolutionary Measures into Conservation Prioritization. *Conserv. Biol.*, 20, 1670–1678.

Rohlf, F.J., Chang, W.S., Sokal, R.R. & Kim, J. (1990). Accuracy of Estimated Phylogenies: Effects of Tree Topology and Evolutionary Model. *Evolution*, 44, 1671–1684.

Ruta, M., Pisani, D., Lloyd, G.T. & Benton, M.J. (2007). A supertree of Temnospondyli: cladogenetic patterns in the most species-rich group of early tetrapods. *Proc. R. Soc. Lond. B Biol. Sci.*, 274, 3087–3095.

Shipley, B. (2000). *Cause and correlation in biology a user's guide to path analysis, structural equations, and causal inference /.*

Shipley, B. (2009). A New Inferential Test for Path Models Based on Directed Acyclic Graphs. *Struct. Equ. Model.*

Shipley, B. (2013). The AIC model selection method applied to path analytic models compared using a d-separation test. *Ecology*, 94, 560–564.

Smith, F.A. & Lyons, S.K. (2013). *Animal body size: linking pattern and process across space, time, and taxonomic group*. University of Chicago Press, Chicago, IL.

Smith, F.A., S. K. Lyons, S. K.M. Ernest, K. E. Jones, D. M. Kaufman, T. Dayan, *et al.* (2003). Body mass of late Quaternary mammals. *Ecology*, 84, 3402 (updated version obtained from senior author).

Stadler, T. (2011). Mammalian phylogeny reveals recent diversification rate shifts. *Proc. Natl. Acad. Sci.*, 108, 6187–6192.

Steel, M. & Mooers, A. (2010). The expected length of pendant and interior edges of a Yule tree. *Appl. Math. Lett.*, 23, 1315–1319.

Tarver, J.E. & Donoghue, P.C.J. (2011). The Trouble with Topology: Phylogenies without Fossils Provide a Revisionist Perspective of Evolutionary History in Topological Analyses of Diversity. *Syst. Biol.*, 60, 700–712.

Thomas, G.H., Hartmann, K., Jetz, W., Joy, J.B., Mimoto, A. & Mooers, A.O. (2013). PASTIS: an R package to facilitate phylogenetic assembly with soft taxonomic inferences. *Methods Ecol. Evol.*, 4, 1011–1017.

Verheyen, W., HULSELMANS, J., WENDELEN, W., LEIRS, H., CORTI, M., BACKELJAU, T., *et al.* (2011). Contribution to the systematics and zoogeography of the East-African Acomys spinosissimus Peters 1852 species complex and the description of two new species (Rodentia: Muridae). *TERMS USE*, 36.

Whitmee, S. & Orme, C.D.L. (2013). Predicting dispersal distance in mammals: a trait-based approach. *J. Anim. Ecol.*, 82, 211–221.

Wilman, H., Belmaker, J., Simpson, J., de la Rosa, C., Rivadeneira, M.M. & Jetz, W. (2014). EltonTraits 1.0: Species-level foraging attributes of the world's birds and mammals. *Ecology*, 95, 2027–2027.

**Table S2.**

**Phylogenetic path model results comparing clades defined by time slices vs. taxa**. Shown are counts of 100 trees where each of 27 models was within 2 CICc units of the best model (and counts passing the d-sep test). The top 3 models are highlighted in gray for each clade delimitation. All models within 2 CICc units are model averaged within each tree; the median of those model averages is in main text Fig. 5 and Fig. S12 for 100 trees.

| | Time slices | | | Taxa | | |
|---|---|---|---|---|---|---|
| **Model name** | **10 Ma** | **30 Ma** | **50 Ma** | **Gen** | **Fam** | **Ord** |
| A1 | 0 (0) | 9 (0) | 25 (1) | 0 (0) | 0 (0) | 0 (0) |
| A1_B1 | 46 (16) | 6 (0) | 0 (0) | 2 (0) | 0 (0) | 0 (0) |
| A1_B1_C1 | 76 (22) | 5 (0) | 0 (0) | 0 (0) | 0 (0) | 0 (0) |
| A1_B1_C3 | 43 (13) | 3 (0) | 0 (0) | 1 (0) | 0 (0) | 0 (0) |
| A1_B3 | 4 (1) | 4 (0) | 59 (2) | 0 (0) | 0 (0) | 0 (0) |
| A1_B3_C1 | 12 (4) | 5 (1) | 14 (0) | 0 (0) | 0 (0) | 0 (0) |
| A1_B3_C3 | 4 (0) | 2 (0) | 8 (0) | 0 (0) | 0 (0) | 0 (0) |
| A1_C1 | 6 (2) | 16 (1) | 2 (0) | 0 (0) | 0 (0) | 0 (0) |
| A1_C3 | 4 (0) | 4 (0) | 5 (0) | 0 (0) | 0 (0) | 0 (0) |
| A3 | 0 (0) | 25 (0) | 52 (0) | 7 (0) | 0 (0) | 0 (0) |
| A3_B1 | 7 (2) | 9 (0) | 0 (0) | 75 (0) | 96 (0) | 0 (0) |
| A3_B1_C1 | 20 (3) | 45 (2) | 3 (0) | 84 (0) | 68 (0) | 0 (0) |
| A3_B1_C3 | 8 (3) | 5 (0) | 0 (0) | 80 (0) | 22 (0) | 0 (0) |
| A3_B3 | 8 (1) | 6 (0) | 45 (0) | 5 (0) | 0 (0) | 0 (0) |
| A3_B3_C1 | 15 (3) | 58 (3) | 23 (0) | 4 (0) | 0 (0) | 0 (0) |
| A3_B3_C3 | 7 (0) | 4 (0) | 2 (0) | 7 (0) | 0 (0) | 0 (0) |
| A3_C1 | 2 (1) | 94 (5) | 35 (1) | 6 (0) | 0 (0) | 3 (2) |
| A3_C3 | 1 (0) | 14 (0) | 7 (0) | 6 (0) | 0 (0) | 0 (0) |
| B1 | 26 (9) | 5 (1) | 1 (0) | 3 (0) | 59 (0) | 0 (0) |
| B1_C1 | 40 (12) | 10 (1) | 0 (0) | 3 (0) | 50 (0) | 0 (0) |
| B1_C3 | 21 (6) | 3 (0) | 0 (0) | 2 (0) | 14 (0) | 0 (0) |
| B3 | 9 (2) | 1 (0) | 53 (1) | 0 (0) | 0 (0) | 0 (0) |
| B3_C1 | 16 (4) | 13 (1) | 35 (1) | 1 (0) | 0 (0) | 0 (0) |
| B3_C3 | 5 (1) | 2 (0) | 9 (0) | 0 (0) | 0 (0) | 0 (0) |
| C1 | 5 (2) | 36 (2) | 11 (1) | 0 (0) | 0 (0) | 88 (63) |
| C3 | 4 (1) | 7 (0) | 3 (0) | 0 (0) | 0 (0) | 50 (30) |
| noEcoVars | 1 (1) | 11 (1) | 20 (0) | 0 (0) | 0 (0) | 0 (0) |
| Percent of models P > 0.05 (d-sep test) | 27.9% | 4.5% | 1.7% | 0.0% | 0.0% | 67.4% |

**Table S3.**

**Summary of path analysis sensitivity tests (Fig. S12).** The number of trees is given (of 1000 tested) where the median path coefficient ± SE did not overlap zero. Having >500 trees with non-zero estimates was viewed as strong evidence (bolded results), ~250-500 trees as marginal, and < 250 trees as zero. Tip-level diversification rates (tip DR) were compared with the node density metric (ND) on the node- and tip-dated full Mammalia trees. Trees were based on all extant non-marine mammals (5,675 species), excluding island endemics (4,553 species), and excluding DNA-lacking and imputed (3,941 species).

| **Test** | **all mammals** | **w/o islands** | **w/o imputed** |
|---|---|---|---|
| Inverse vagility ~ tip rate mean, 10 Ma | | | |
| tip DR, node-dated | **854** | **701** | **555** |
| tip DR, tip-dated | 457 | 442 | 271 |
| ND, node-dated | **959** | **929** | **998** |
| ND, tip-dated | **862** | **675** | **979** |
| Positive diurnality ~ tip rate mean, 10 Ma | | | |
| tip DR, node-dated | **746** | 497 | **740** |
| tip DR, tip-dated | **573** | 417 | **548** |
| ND, node-dated | **886** | **762** | **714** |
| ND, tip-dated | 316 | 146 | 133 |
| Positive latitude ~ tip rate mean, all slices | | | |
| tip DR, node-dated | **591, 722, 572** | **756, 930**, 471 | 475, **644**, 255 |
| tip DR, tip-dated | 459, 423, **698** | **603, 626, 703** | 264, 228, 366 |
| ND, node-dated | 47, 2, 151 | 69, 45, 187 | 454, 0, 146 |
| ND, tip-dated | 20, 2, **525** | 74, 65, **608** | 79, 4, **521** |

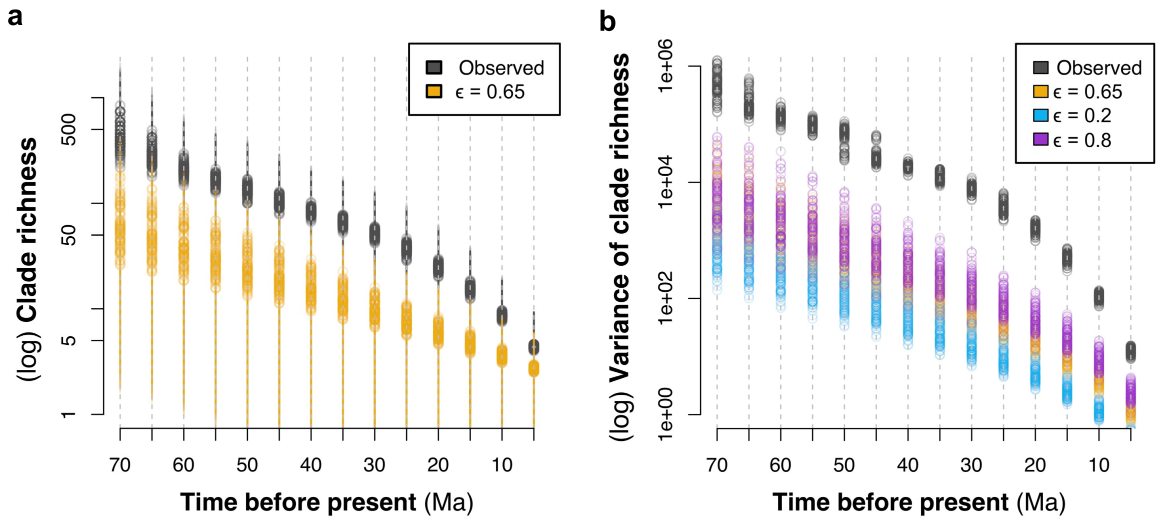

**Fig. S6**

**Characterizing tree-wide time-sliced clades from 5-70 Ma. (a)** Variation in the species richness of time-sliced clades delimited every 5 Ma from 5-70 Ma is shown as log clade richness and (**b**) log variance of clade richness, comparing values from observed mammal clades (grey, 100 trees drawn from the node-dated credible set of completed trees) relative to simulated clades from trees using different extinction fractions, ε (colors; rate-constant birth-death [RCBD] simulations of 100 trees). Error bars in **part a** are 95% confidence intervals (CIs). Observed mammal clades consistently have higher and more variable species richness than expected under the RCBD simulations.

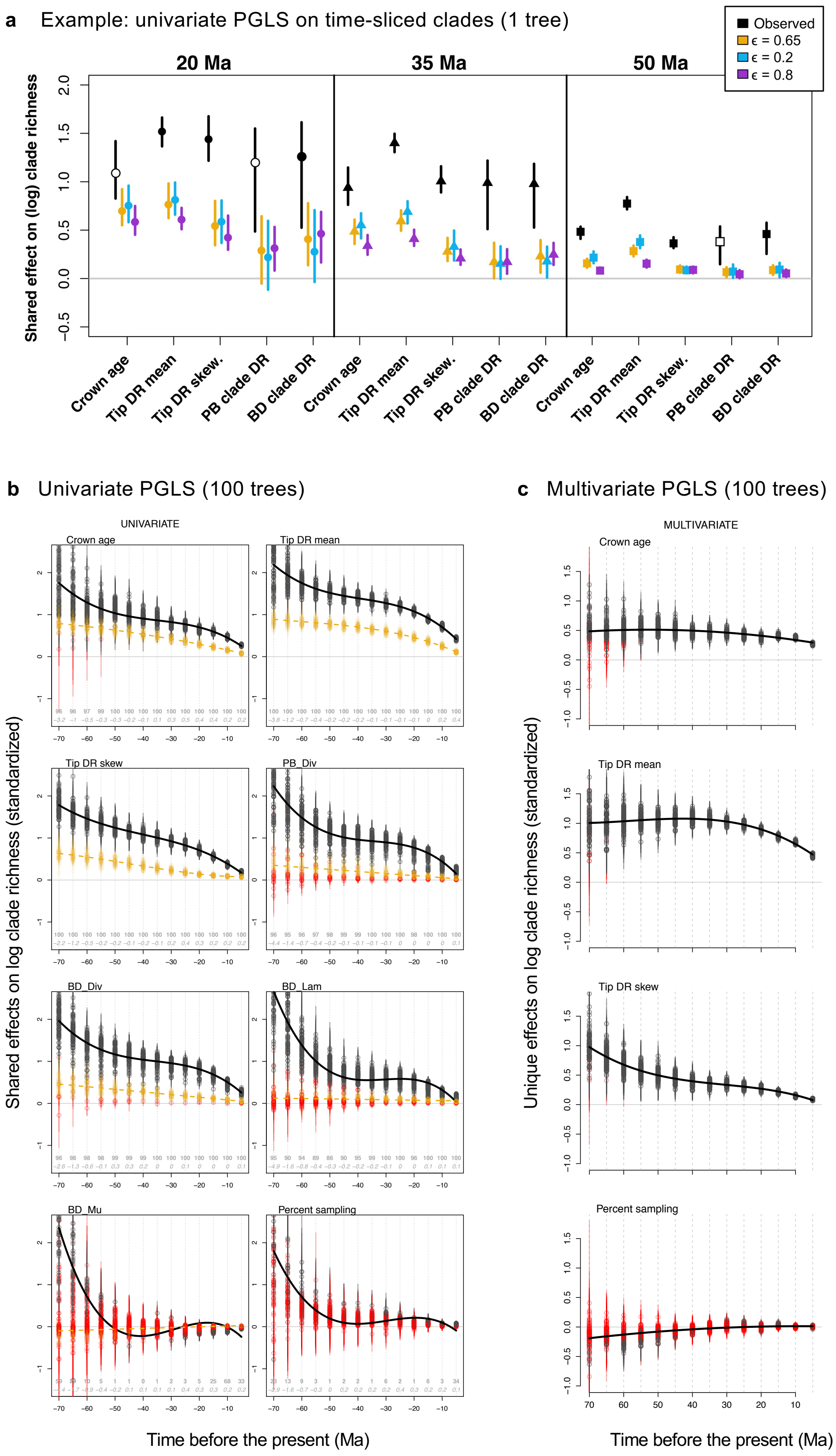

**Fig. S7**

**Tests of factors explaining log clade species richness.** (**a**) Example of univariate phylogenetic generalized least squares (PGLS) regression performed as log clade richness ~ predictor across five clade-level metrics: clade crown age, tip DR harmonic mean, tip DR skewness, pure-birth (PB) diversification rate, and birth-death (BD) net diversification rate. Predictors are standardized as mean centered and standard deviation scaled. Clades were delimited at 20-, 35-, and 50-Ma time slices for 1 tree as an example, showing shared effects and 95% CIs (black) against rate-constant birth-death (RCBD) simulations using different extinction fractions, ε (colors). Open symbols indicate overlap in the 95% CIs while closed symbols are significantly different. (**b**) Full univariate results from time-sliced clades across 100 trees (black) as compared to RCBD simulations of ε = 0.65 (yellow). Additional predictors include: birth-death speciation rate (BD_Lam), birth-death extinction rate (BD_Mu), and the percentage of DNA-sampled species (Percent sampling). Bold numbers per time slice are the percentage of significant results of 100 trees, and italic numbers are the mean values of Pagel's lambda estimated during PGLS. (**c**) Full multivariate results for the most predictive variables, showing their unique effects across time-sliced clades delimited on 100 mammal trees and including percent sampling. Red symbols are non-significant effects in which the 95% confidence interval overlaps zero.

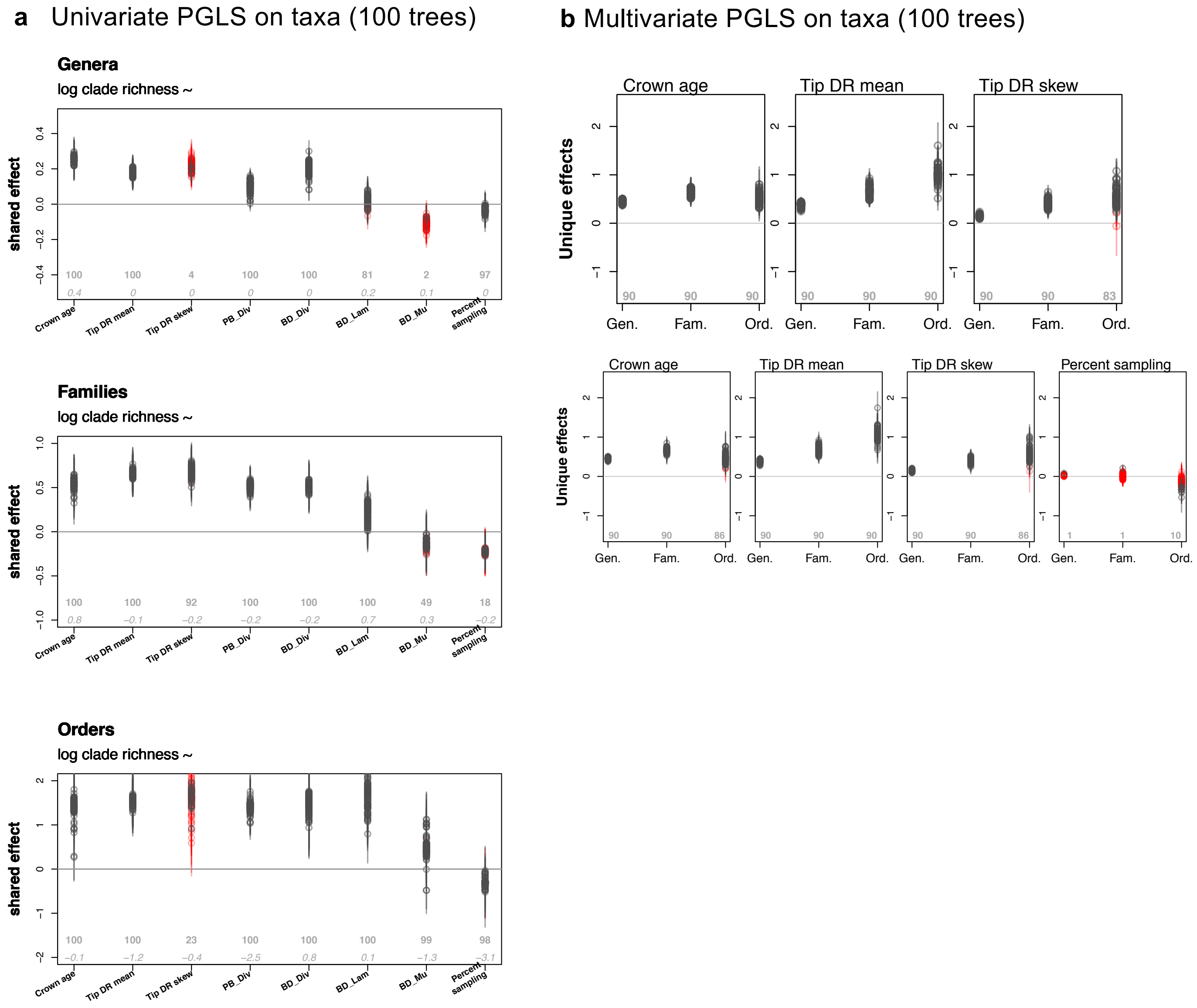

**Fig. S8**

**Taxon-based clade richness predictors.** Analogous tests of log clade richness ~ predictors for genera, families, and orders of mammals, conducting PGLS analyses in (**a**) univariate and (**b**) multivariate contexts across 100 mammal trees. Gray circles represent significant terms in the model, while red triangles have $P > 0.05$. Only clades with ≥4 species were examined for genera (n=385), families (n=102), and orders (n=22). **Part b** includes a lower panel in which percent sampling is included in the multivariate model to test for any bias from using taxonomically completed trees. Clade-level predictors shown are crown age, the harmonic mean of tip DR values (tip DR mean), the skewness of tip DR values (tip DR skew), pure-birth diversification rate (PB_Div), birth-death diversification rate (BD_Div), birth-death speciation rate (BD_Lam), birth-death extinction rate (BD_Mu), and the percentage of DNA-sampled species (Percent sampling).

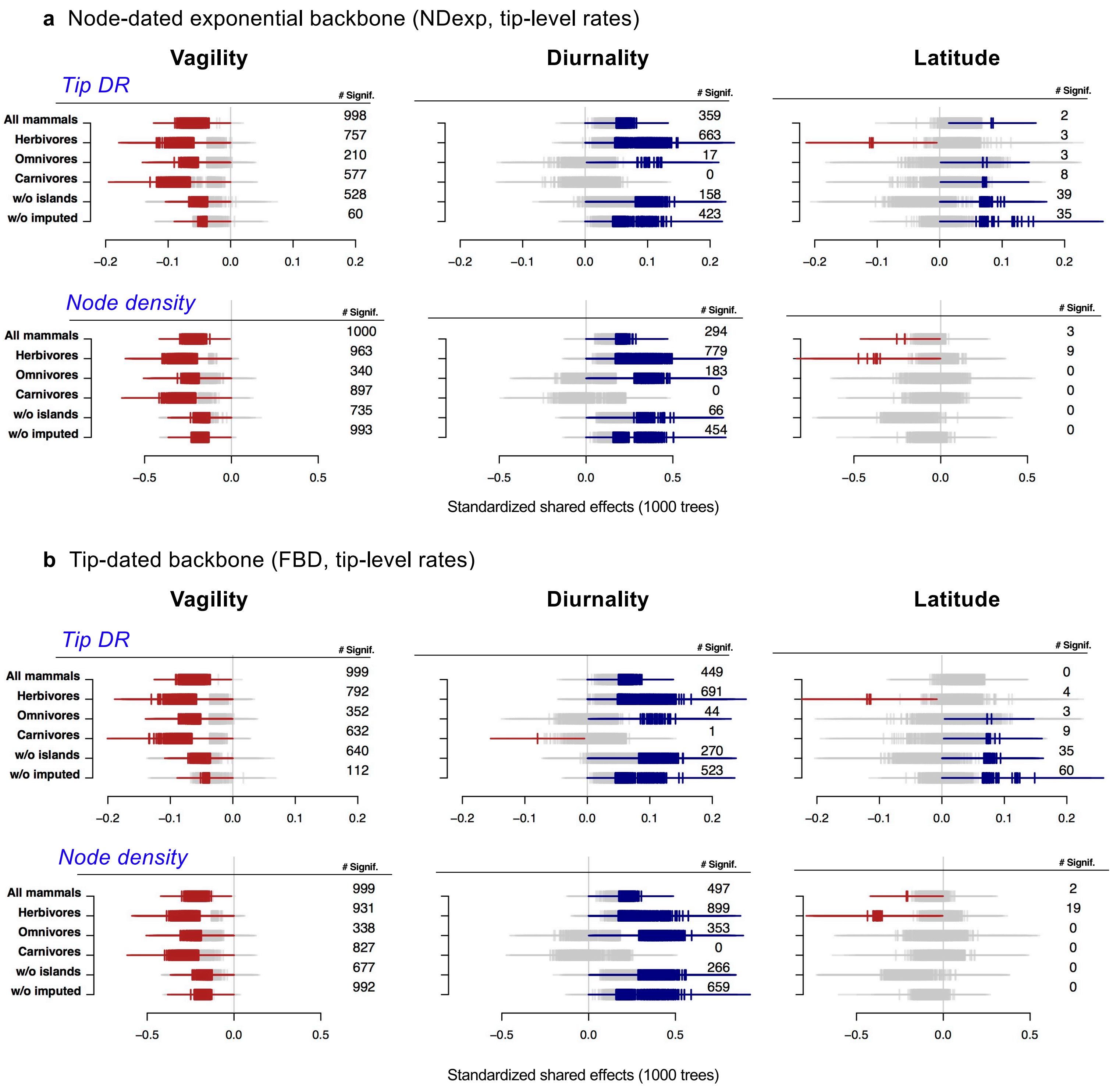

**Fig. S9**

**Tip-level sensitivity analyses of univariate PGLS models (see Fig. 5a).** Analyses of trait ~ rate models are repeated across 1,000 trees from the completed tree credible sets for (**a**) the node-dated backbone, and (**b**) the tip-dated backbone, in each case comparing standardized shared effects of traits upon tip DR and node density metrics. The following data subsets were examined: all mammals (extant and non-marine for all trait analyses; $N = 5{,}675$), herbivores only ($N = 1{,}637$), omnivores only ($N = 1{,}852$), carnivores only ($N = 1{,}565$), excluding island endemics (strict classification, $N = 761$), and excluding taxonomically imputed species in the completed trees ($N = 1{,}813$). The right-side margin shows totals of non-zero estimates with $P < 0.05$, either positive (blue shades) or negative (red shades) out of the 1,000 trees analyzed (gray bars denote effects of $P > 0.05$).

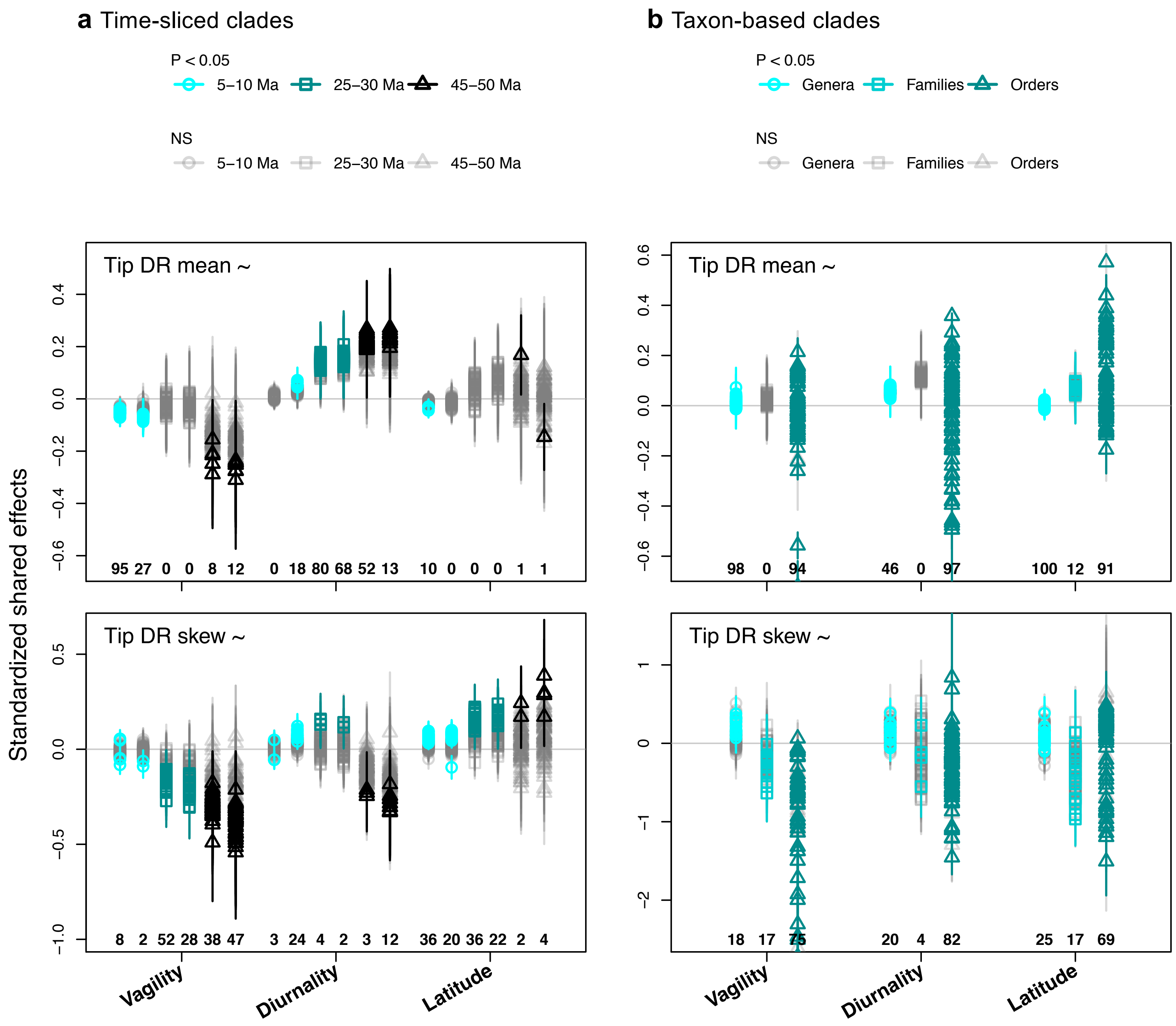

**Fig. S10**

**Clade-level univariate PGLS analyses of ecological factors on tip DR mean and skew.** Results across 100 mammal trees are compared using (**a**) time-sliced clades delimited at representative slices of 5, 10, 25, 30, 45, and 50 Ma (symbols in legend at top); and (**b**) taxon-based (named) clades of mammal genera, families, and orders. The number of trees for which a given univariate comparison was significant is displayed below each set of standardized shared effects.

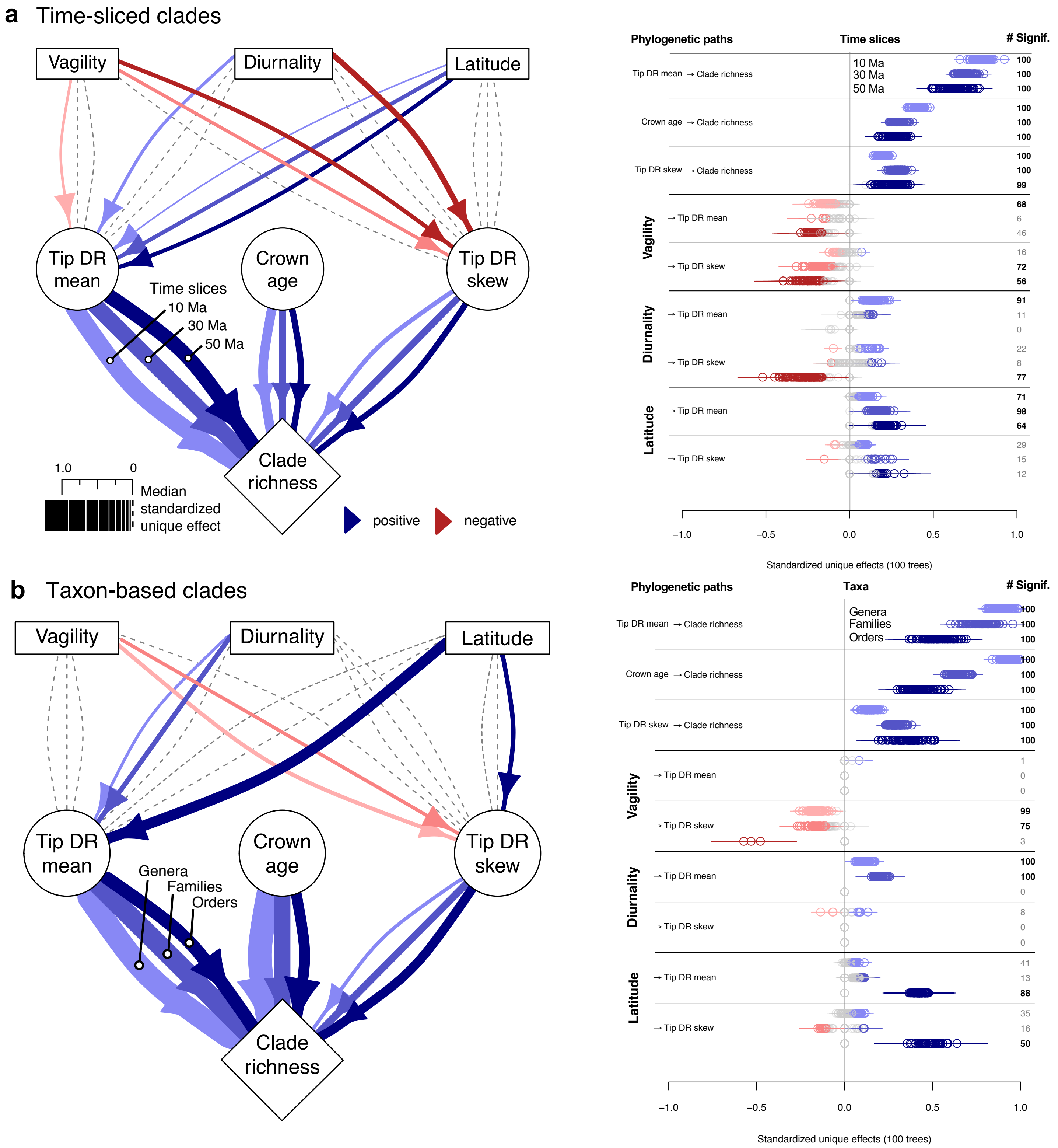

**Fig. S11**

**Expanded display of Fig. 5 path analyses comparing time-sliced clades vs. taxa.** (**a**) Clades defined by time slices at 10-, 30-, and 50-Ma intervals (light to dark colors). (left) Path thickness and directionality denotes median coefficients of model-averaged analyses on standardized data and 100 trees. (right) Per-estimate uncertainty across analyses and taxa (slope ± SE, 100 trees). (**b**) The same models run upon clades delimited as genera, families, and orders (light to dark colors). The right-side margin shows totals of non-zero estimates, either positive (blue shades) or negative (red shades), and bolds the totals of paths present in >50 trees (grey denotes majority overlap with zero, which are dashed-line paths on the left-side).

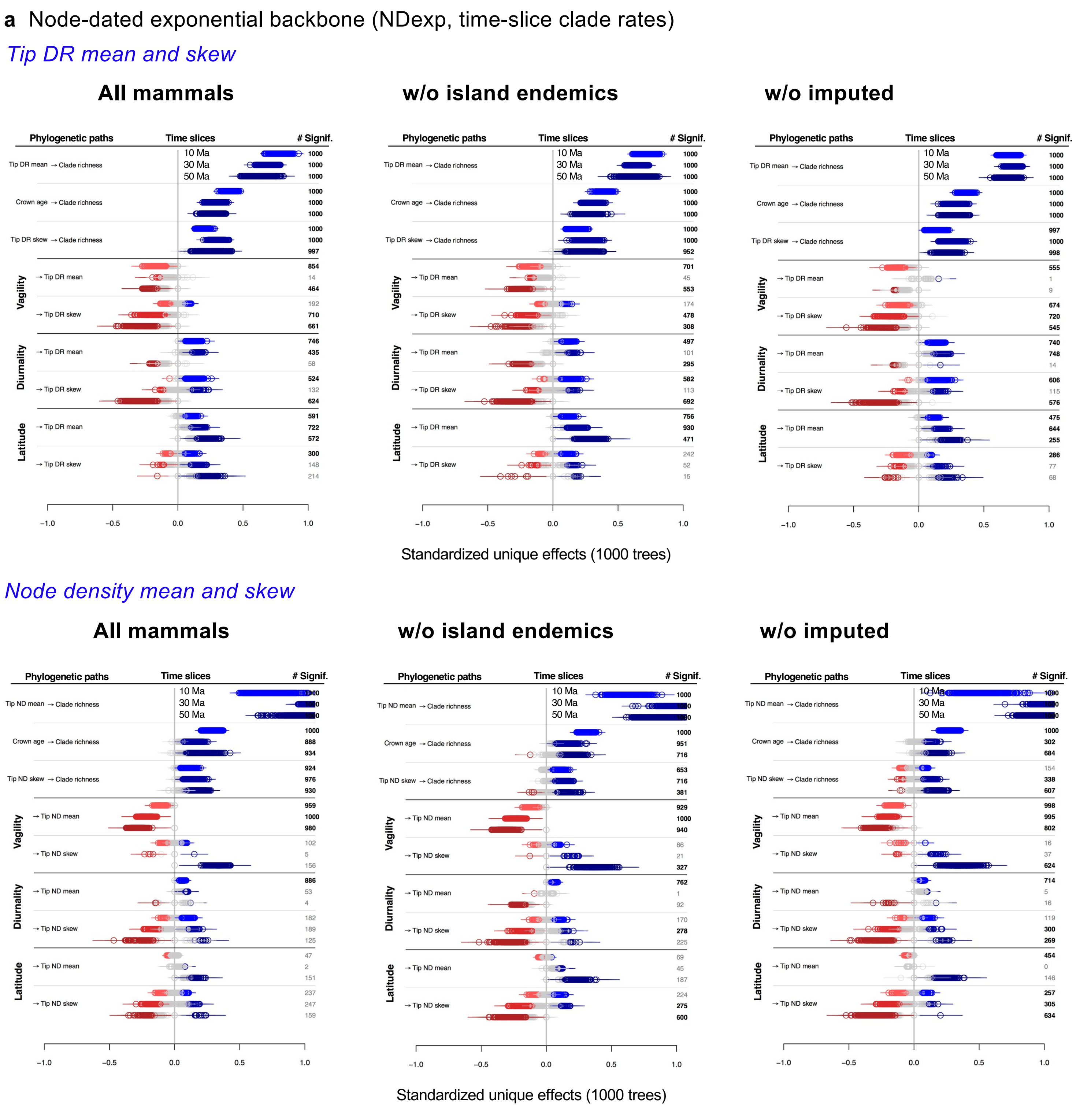

**Fig. S12**

**Clade-level sensitivity analyses of multivariate phylogenetic path models (see Fig. 5b).** Comparison of path coefficients on time-sliced clades at 10, 30, and 50 Ma delimited across 1,000 trees from the taxonomically completed credible sets for (**a**) the node-dated backbone, and (**b**) the tip-dated backbone. Path models were run on the following data subsets: all mammals (extant and non-marine for all trait analyses; $N = 5{,}675$), excluding island endemics (strict classification, $N = 761$), and excluding taxonomically imputed species in the completed trees ($N = 1{,}813$). Median coefficients ± SE are shown for model-averaged runs using standardized data. The right-side margin shows totals of non-zero estimates with $P < 0.05$, either positive (blue shades) or inverse (red shades), and bold totals of paths present in >500 of the 1,000 trees analyzed (gray denotes $P > 0.05$).

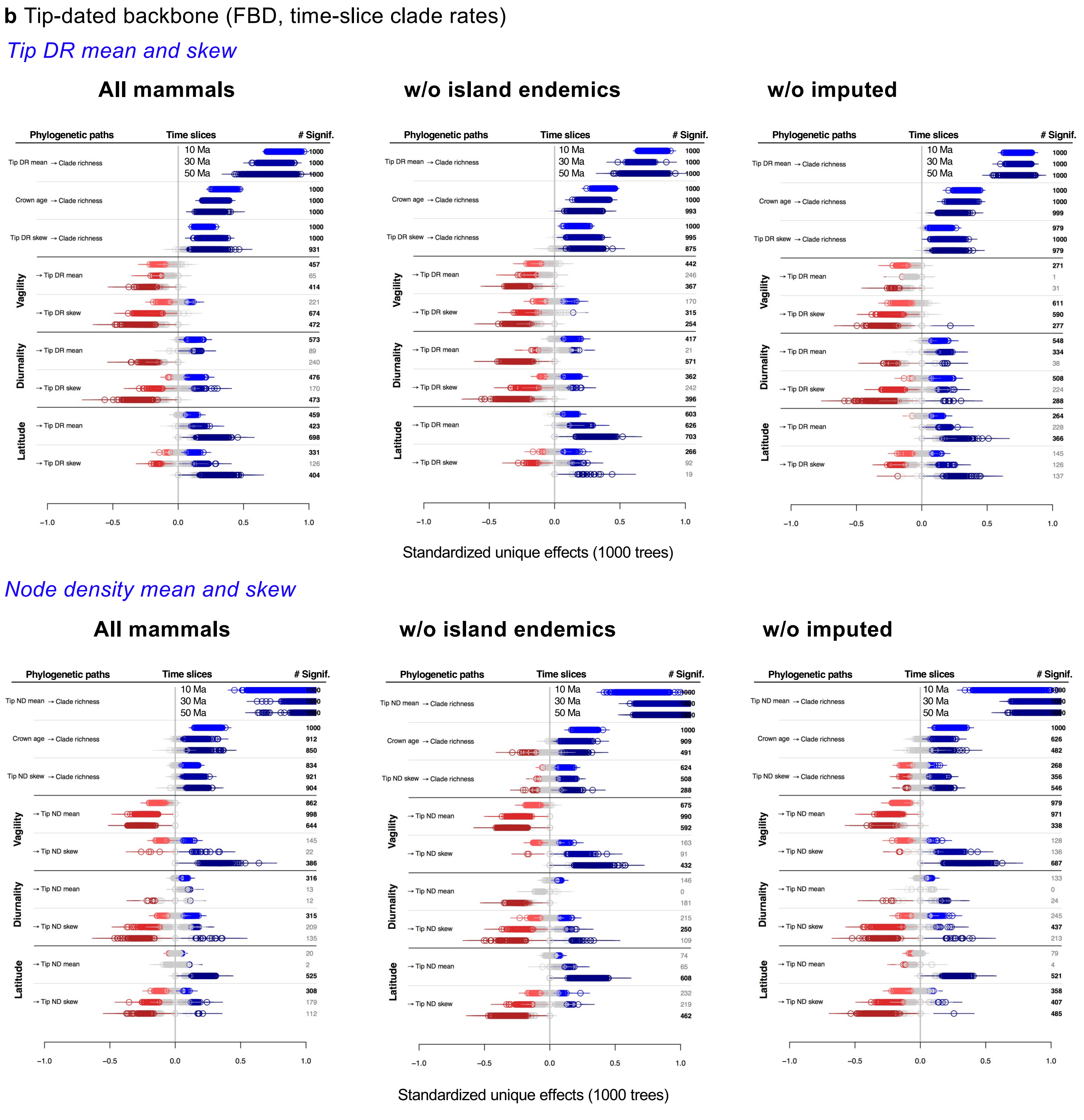

**Fig. S12 (continued)**

**Clade-level sensitivity analyses of multivariate phylogenetic path models.** See previous legend.
